# Hydrogel confinement of the secreted nascent matrix imposes solid stress and mediates pathology of chondrocytes

**DOI:** 10.64898/2026.10.19.760217

**Authors:** Jake Song, Sarah Loveland, Fan Yang, Nidhi Bhutani, Marc Levenston, Ovijit Chaudhuri

**Affiliations:** Department of Mechanical Engineering, Stanford University, Stanford, CA, 94305; Department of Chemistry, Stanford University, Stanford, CA, 94305; Department of Bioengineering, Stanford University, Stanford, CA, 94305; Department of Orthopedic Surgery, Stanford University, Stanford, CA, 94305

## Abstract

Mechanical confinement of cells in 3D hydrogel scaffolds mediates cell fate and activity across a wide range of biological processes. Many cells secrete a nascent matrix within the hydrogel niche, which provides important biological signals to the cells but also obstructs direct cell-hydrogel interactions, making it unclear how the cells sense hydrogel confinement and carry out mechanotransduction. Here, we show that chondrocytes sense the solid stress arising from the accumulation of the nascent matrix in the hydrogel niche, which is mediated by hydrogel viscoelasticity and plasticity. We show that solid stresses elicit a wide range of pathophysiological responses, most notably an inflammatory response associated with cartilage osteoarthritis and injury, alongside a striking progenitor-like migratory response where the chondrocytes leverage mitosis to breach their nascent matrix and migrate to escape confinement. Our findings establish the role of the nascent matrix as an important transducer of mechanical confinement and illustrate how emergent solid stresses can drive chondrocyte pathology.

## Main

In disease modeling and tissue engineering applications, cells are routinely encapsulated in 3D hydrogel scaffolds to mimic their native tissue microenvironments – a process which inevitably leads to their mechanical confinement. Confinement arises as the hydrogels are nanoporous and physically surround the cells, thereby restricting basic cellular processes such as growth, proliferation, and migration.^1,2^ The extent of confinement is governed by material properties of hydrogels that are often tuned to mimic those of native tissues, including not only static properties like stiffness or elasticity but also dynamic properties that enable remodeling such as viscoelasticity (with time),^3,4^ plasticity (with strain),^5,6^ and degradability (with proteolysis)^7^. Together, these material properties mediate the mechanical confinement experienced by the cells, and direct cell fate across a range of developmental and disease contexts.^1^

Understanding the mechanisms by which cells sense mechanical confinement can provide insight into cell-matrix interactions that underlie development and disease *in vivo*, and can guide the rational design of cell scaffold materials. Canonically, this process is assumed to be mediated by direct cellular interactions with hydrogel scaffolds, for instance by integrin-mediated tethering via RGD functional groups, or by direct physical restriction of volumetric growth.^1,8^ However, recent studies have shown that cells such as mesenchymal stromal cells (MSCs) and chondrocytes also secrete their own “nascent” matrix that accumulates in the hydrogel cavity, which can physically displace the hydrogel and prevent direct cell-hydrogel interactions.^9–11^ In particular, studies on MSCs have shown that the nascent matrix provides important biological ligands for cell spreading and differentiation, yet cells remained sensitive to the confining properties of the hydrogel such as stiffness, viscoelasticity, and degradability.^9,11^ How matrix-secreting cells sense the mechanical confinement of their microenvironment, and the role played by the nascent matrix in this mechanosensory process, remain poorly understood (Fig. 1A).

**Figure 1.**
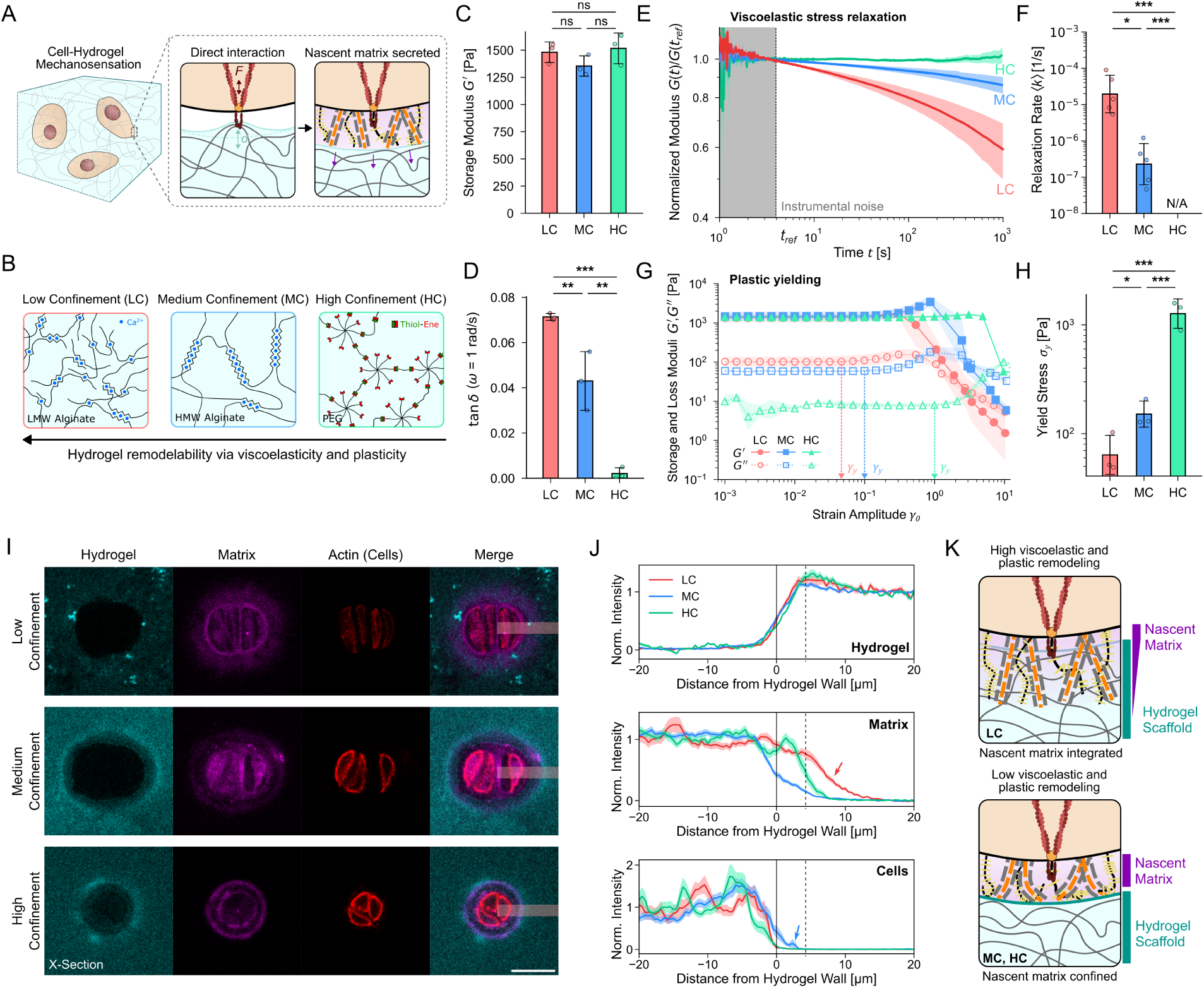
Hydrogel remodelability via viscoelasticity and plasticity mediates confinement of the chondrocyte nascent matrix. **A)** Matrix-secreting cells such as chondrocytes secrete their nascent matrix within the hydrogel niche, which can obstruct direct cell-hydrogel interactions. Here, we study how such matrix-secreting cells sense hydrogel mechanical confinement. **B)** Schematic illustration of the low confinement (LC), medium confinement (MC), and high confinement (HC) hydrogel scaffolds used in the study, consisting of low molecular weight (LMW) alginate, high molecular weight (HMW) alginate, and thiol-ene cross-linked poly(ethylene glycol) (PEG). **C, D)** Storage modulus *G*′ and tan *δ* of the three hydrogel systems at *ω* = 1 rad/s. **E)** Averaged traces of the relaxation modulus *G*(*t*) (normalized to a reference time of ∼ 4 seconds) for the three hydrogel systems in response to a linear step strain. **F)** Mean relaxation rate, 〈*k*〉, obtained by fitting the stress relaxation traces in panel E with the Kohlrausch-Williams-Watts function (see Fig. S1 for fits). N = 5, 5, and 3 for LC, MC, and HC. **G)** Averaged storage modulus *G*′ and loss modulus *G*′′ obtained from amplitude sweeps. The yield strain, *γ_y_*, is defined as the point at which *G*′′ begins to increase due to unrecoverable plastic deformations.^70^ Shown on the plot are averages of *γ_y_* for the three systems (see Fig. S2 for quantification protocol). N = 3 for all systems. **H)** Yield stress *σ_y_*, defined as the corresponding oscillatory stress at *γ_y_*. **I)** Representative images of chondrocytes in the different hydrogels at day 7 – note that all chondrocytes are seeded as single cells, and proliferate into multi-cellular clusters over time. Matrix staining is done by metabolically labeling nascent proteins with azidohomoalanine (AHA), and visualized with DBCO-555. Scale bars 25 µm. **J)** Averaged intensity linescans of the hydrogel, matrix, and cell channels centered on the hydrogel cavity wall (shaded area sampled in panel I). Dashed lines mark the end point of the cavity wall (represented by the maximum fluorescence intensity in the hydrogel channel). Arrow in the Matrix panel highlights the significant matrix penetration into the hydrogel under low confinement. Arrow in the Cell panel highlights the proximity of cells to the hydrogel wall. See Methods for normalization protocols. N = 16, 28, and 15 for LC, MC, and HC. **K)** Summary illustration demonstrating nascent matrix integration in low confinement hydrogels, and nascent matrix confinement in medium and high confinement hydrogels.

Here, we demonstrate that the material properties of the hydrogel such as stiffness, viscoelasticity, and plasticity can control the confinement of the nascent matrix within the hydrogel cavity, thus imposing a compressive solid stress on the resident cells. We demonstrate this effect on matrix-secreting chondrocytes and illustrate how confinement-mediated solid stresses induce an inflammatory response alongside a surprising migratory response reminiscent of those reported in chondroprogenitors that emerge during cartilage mechanical injury and osteoarthritis.

### Hydrogel remodelability mediates mechanical confinement of encapsulated chondrocytes

To investigate this problem, we first encapsulated matrix-secreting juvenile bovine chondrocytes in hydrogels with similar elastic moduli but varying inelastic properties that enable the hydrogels to be remodeled with time (viscoelasticity) and deformation (plasticity). To engineer these properties, we used low molecular weight and high molecular weight alginate hydrogels, which exhibit high and low levels of viscoelastic and plastic responses, respectively^3,5,12^. Additionally, we used cross-linked poly(ethylene glycol) hydrogels as a primarily elastic system with negligible viscoelastic and plastic responses (Fig. 1B).^13^ Since viscoelasticity and plasticity enable hydrogel remodeling and reduce effective confinement experienced by the cells, we henceforth refer to these hydrogels as low confinement (LC), medium confinement (MC), and high confinement (HC), respectively. These hydrogels lack adhesive ligands (as adhesive ligands often promote dedifferentiation of chondrocytes^14^) or biodegradable elements, allowing us to study the effect of mechanical confinement in isolation.

We performed systematic rheological characterizations of the three hydrogel systems to validate these mechanical properties. Rheological characterizations show that the three hydrogel systems exhibit similar elastic moduli (Fig. 1C), but varying levels of viscoelastic dissipation (Fig. 1D), and stress relaxation rates (Fig. 1E, 1F, S1). Low confinement hydrogels exhibited the highest level of viscoelastic dissipation and stress relaxation rates, followed by medium confinement and by high confinement. Similar trends were observed with yield stress (the stress at which irreversible plastic deformations begin to accrue – Fig. 1G, 1H, S2) and ultimate failure stress (the stress at which the material macroscopically fails – Fig. S3). Rheological characterizations show that low confinement hydrogels exhibit the earliest onset of irreversible plastic deformations and the lowest ultimate failure stresses, followed by medium confinement and by high confinement. Altogether, we designed hydrogels with similar elasticity and different extents of viscoelastic and plastic remodelability, which thus confer low, medium, and high confinement onto encapsulated cells (Fig. 1B). It is noted that our experimental setup cannot rigorously distinguish between confinement effects on cells arising from viscoelastic versus plastic remodeling – these properties are often coupled in non-covalently cross-linked gels and are difficult to separate^15^ – but we emphasize that both of these properties will facilitate cell-mediated remodeling of the scaffold, and thus control the confinement experienced by the cells.

**Figure 2.**
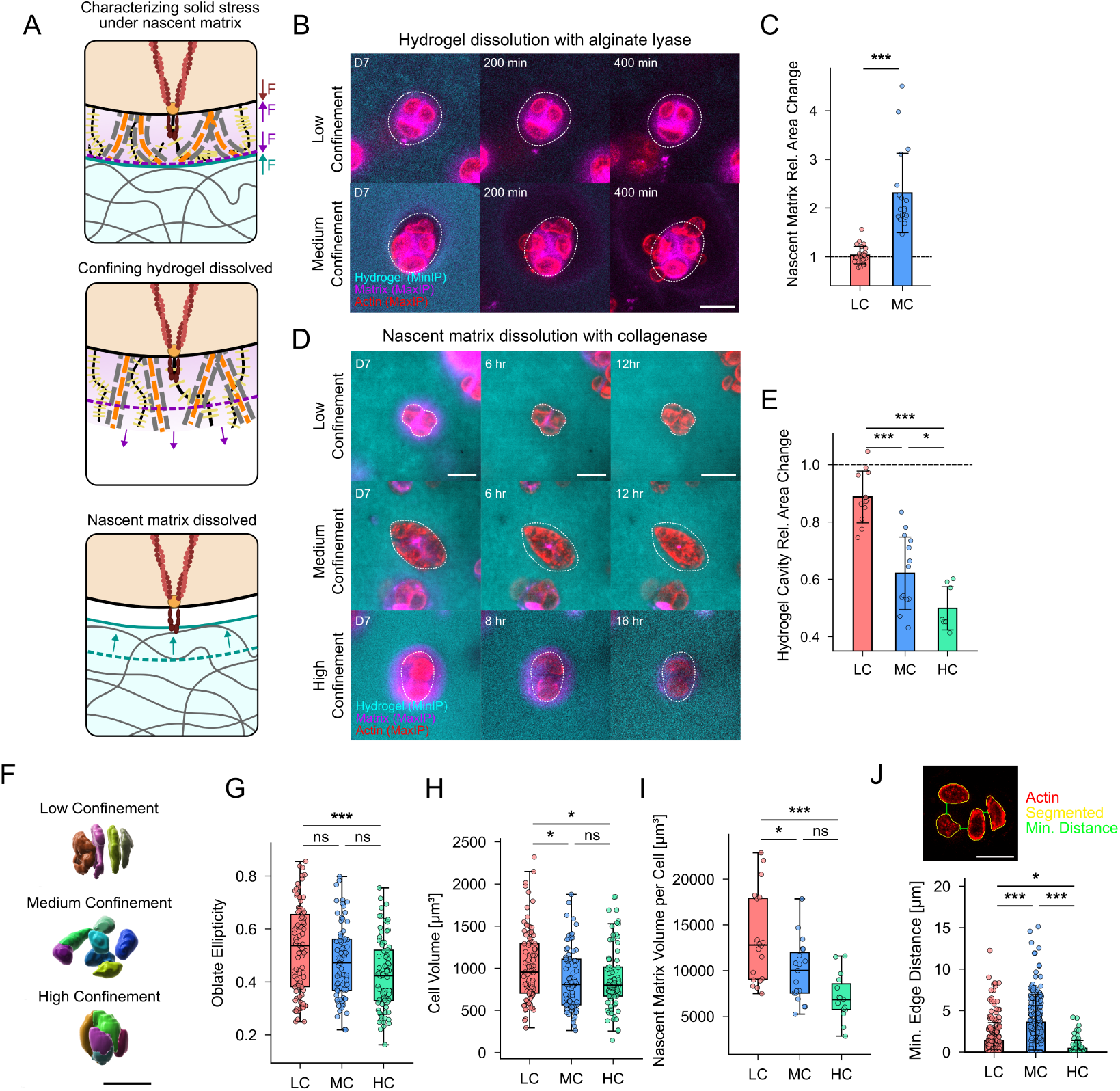
Nascent matrix confinement imposes solid stress on chondrocyte clusters and alters cell morphology. **A)** Hypothesis for investigating whether nascent matrix confinement leads to solid stress. Under solid stress, dissolving the confining hydrogel should lead to an expansion of the nascent matrix, and dissolving the nascent matrix should lead to a contraction of the hydrogel cavity. **B)** Representative time-lapse images of cell clusters in low and medium confinement hydrogels during dissolution with alginate lyase. Dashed lines indicate initial area of the nascent matrix. See Fig. S7 for additional examples, and S8 for post-mortem images after hydrogel dissolution. **C)** Area change in the nascent matrix after hydrogel dissolution relative to before. **D)** Representative time-lapse images of cell clusters in low, medium, and high confinement hydrogels during matrix dissolution with collagenase. **E)** Relative hydrogel cavity area change after matrix dissolution compared to before. **F)** Representative 3D segmentation of cell clusters in low, medium, and high confinement hydrogels. Scale bar 25 µm. **G, H, I)** Oblate ellipticity, cell volume, and cluster nascent matrix volume (normalized to number of cells within the cluster) in low, medium, and high confinement hydrogels. **J)** Minimum edge-to-edge distance between cells within clusters in low, medium, and high confinement hydrogels. Note that this characterization was performed on 2D cross-sectional images. Scale bars 25 µm.

### Hydrogel remodeling mediates confinement of the nascent matrix in the hydrogel cavity

Using hydrogels with tunable viscoelastic and plastic remodeling, we studied the spatiotemporal evolution of chondrocytes and their secreted nascent matrix within fluorescently labeled hydrogels. All chondrocytes were filtered and seeded as single cells, but proliferated into multi-cellular clusters by day 7 (Fig. 1I, S4). To visualize the nascent matrix, we metabolically labeled nascent proteins including collagens with an azide-modified methionine analog azidohomoalanine (AHA) (represented by default in figure panels) or sugar-containing glycosaminoglycans with azide-modified sugar analogs N-azidoacetylmannosamine (ManNAz) and N-azidoacetylgalactosamine (GalNAz).^10,16^

By staining the nascent matrix, we found striking differences in the manner in which the nascent matrix interfaces with the hydrogels: whereas the nascent matrix penetrated into hydrogels in low confinement hydrogels, the nascent matrix instead became confined within the hydrogel cavity in medium and high confinement hydrogels (Fig. 1I). Intensity linescans of the hydrogel, matrix, and cell channels showed that while the hydrogel is displaced across all three confinement conditions – evidenced by the densification of the intensity signal at the edge of the cavity – only low confinement facilitates significant penetration of the nascent matrix proteins through this densified hydrogel interface (Fig. 1J). A similar linescan analysis of nascent matrix glycans revealed a similar, but less dramatic, difference in matrix penetration across the three hydrogels (Fig. S6), which suggests that the more flexible glycan biopolymers exhibit greater interpenetration with the hydrogels, consistent with a previous study on glycan penetration^10^. In summary, we found that hydrogel confinement caused accumulation of the nascent matrix at the cell-hydrogel interface, but the alleviation of this confinement via viscoelastic and plastic remodeling instead resulted in nascent matrix penetration into the hydrogels (Fig. 1K).

### Nascent matrix confinement imposes solid stress on chondrocyte clusters, altering cell morphology

Given the observations of nascent matrix confinement in hydrogels with low or minimal viscoelastic and plastic remodeling, we hypothesized that the continued accrual of the nascent matrix under confinement could result in a build-up of pressure within the hydrogel niche, and thus impose compressive “solid stresses” on cells.^17^ If the matrix is pressurized from solid stresses, we reasoned that the dissolution of the hydrogel would lead to an expansion of the nascent matrix, and the dissolution of the nascent matrix would lead to a contraction of the hydrogel (Fig. 2A). First, dissolution of the alginate polymer in low confinement and medium confinement hydrogels using alginate lyase showed that, while the nascent matrix area remained unchanged when dissolving low confinement hydrogels, the matrix area increased significantly when dissolving medium confinement hydrogels (Fig. 2B, 2C, Movie 1); this observation was consistent for nascent matrix proteins as well as nascent matrix glycans (Fig. S7, Movie 2).

Interestingly, we also observed cells escaping the nascent matrix in medium confinement hydrogels during hydrogel dissolution. Next, dissolution of the nascent matrix in all three hydrogels using collagenase revealed that the hydrogel area remained mostly unchanged in low confinement hydrogels but shrank significantly in medium and high confinement hydrogels (Fig. 2D, 2E, Movie 3). Together, these findings indicate that nascent matrix confinement, which is mediated by mechanical properties of the hydrogel scaffolds, imposes solid stress on the cell clusters.

Based on cavity area changes in Fig. 2E, it is possible to crudely estimate the solid stress imposed by nascent matrix confinement on the cells in the hydrogels. For a spherical cavity growing in a neo-Hookean elastic medium, the cavity pressure can be derived as 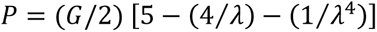 where 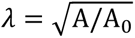 is the circumferential stretch of the cavity, and A is the inflated area and A_0_ is the resting area, respectively.^18^ Substituting the shear moduli *G* (Fig. 1C) and area change (Fig. 2E) into this relation, we obtain estimated cavity pressures of *P* ∼ 300, 1000, and 1500 Pa for low, medium, and high confinement hydrogels, respectively. These values should only be considered as estimates, as they neglect contributions from cavity morphology, plasticity and fracture, viscoelasticity, and strain stiffening. Additionally, stress may not directly be absorbed by the cells, as the nascent matrix can be stiff and load-bearing (on the order of 10 kPa in elastic modulus^10^). Nevertheless, considering that stresses on the order of 100-1000 Pa are sufficient to induce significant changes to cell fate, including migration, blebbing, or death,^19–21^ these estimated values indicate that nascent matrix confinement in our hydrogels can provide relevant magnitudes of solid stresses to alter cell fate and induce changes to cell phenotype.

We then investigated the morphological consequences of solid stress on the cells. Segmentation of the cells within the clusters in 3D showed that cells in low confinement hydrogels were arranged like stacked discoids (Fig. 2F), thus exhibiting high oblate ellipticity (Fig. 2G). These morphological features resemble those observed in proliferating cells within the cartilage. On the other hand, cells in medium and high confinement hydrogels were rounder, evidenced by reduced ellipticity (Fig. 2G), and smaller, evidenced by lower cell volumes (Fig. 2H). The cells also secreted a smaller volume of nascent matrix (Fig. 2I). Cells in high confinement hydrogels were also tightly packed together, indicated by the low cell-cell distances within the clusters (Fig. 2J). These observations show that solid stress imposes a morphological change in chondrocytes, including cell rounding, volume restriction, and cluster compaction.

As solid stress arises from the cavity pressure caused by the elastic resistance of the hydrogels to the volume increase from nascent matrix secretions, we hypothesized that increasing hydrogel stiffness should impose greater confinement-induced solid stress on the cells and affect cell morphology, in a similar vein to reducing viscoelastic and plastic remodeling, and vice versa. Previous works have shown that increasing hydrogel stiffness reduces the volume of the nascent matrix,^9,10^ and while this has been implicated in affecting MSC differentiation,^9^ the mechanical consequence of this nascent matrix volume restriction has remained unknown. To test this, we varied the shear modulus of the low confinement and medium confinement alginate hydrogels by almost an order of magnitude by tuning the total concentration of polymer and calcium cross-links (Fig. S9). While cell morphology was independent of elasticity in low confinement hydrogels, the morphology was highly dependent on shear moduli in medium confinement hydrogels. Compared to a baseline behavior at ∼ 1400 Pa, softer formulations of ∼ 900 Pa promoted discoidal morphologies found in low confinement, whereas stiffer formulations of ∼ 2700 Pa promoted greater cluster area, cell-cell distance, and densification of the hydrogel. At an even stiffer formulation of ∼ 6900 Pa, cell clusters began shrinking into smaller, rounder, and compacted clusters reminiscent of those observed in high confinement. These findings provide further evidence that material properties of hydrogel scaffolds, such as stiffness, viscoelasticity, and plasticity, mediate solid stress imposed on cells via nascent matrix confinement.

### Medium confinement promotes directional growth of chondrocyte clusters

We next performed a temporal investigation of the cell clusters via longitudinal time-lapse imaging (Fig. 3A). Nuclei segmentation showed that, while cells in low and medium confinement hydrogels proliferated similarly, cells in high confinement hydrogels showed limited proliferation evidenced by a stagnant growth in nuclei count (Fig. 3B). This was independently validated by dsDNA quantifications in the hydrogels at day 7 (Fig. 3C), and indeed, dead nuclei fragments were visible throughout the images (Fig. S10). In a similar vein, reverse transcription quantitative polymerase chain reaction (RT-qPCR) of the expression of canonical senescence marker *p16* showed significant upregulation in cells in high confinement hydrogels compared to those in low and medium confinement hydrogels, supporting the interpretation of limited proliferation under high confinement (Fig. 3D). These observations are unusual, as cells from hyaline juvenile cartilage normally exhibit a proliferation program^22^, and suggest the role of hydrogel confinement in arresting this program. In support of this interpretation, multiple cells were observed to be arrested in telophase, unable to pinch off their cortex into two separate bodies and thus undergo cytokinesis (Fig. 3E, S10). As telophase and cytokinesis stages of mitosis generate the largest amount of protrusive forces on the surrounding microenvironment,^23^ cell cycle arrest at cytokinesis suggests the inability of the cell to surmount the confining stresses of the hydrogel scaffold. Altogether, these results show that high solid stresses promoted cell cycle arrest and death.

**Figure 3.**
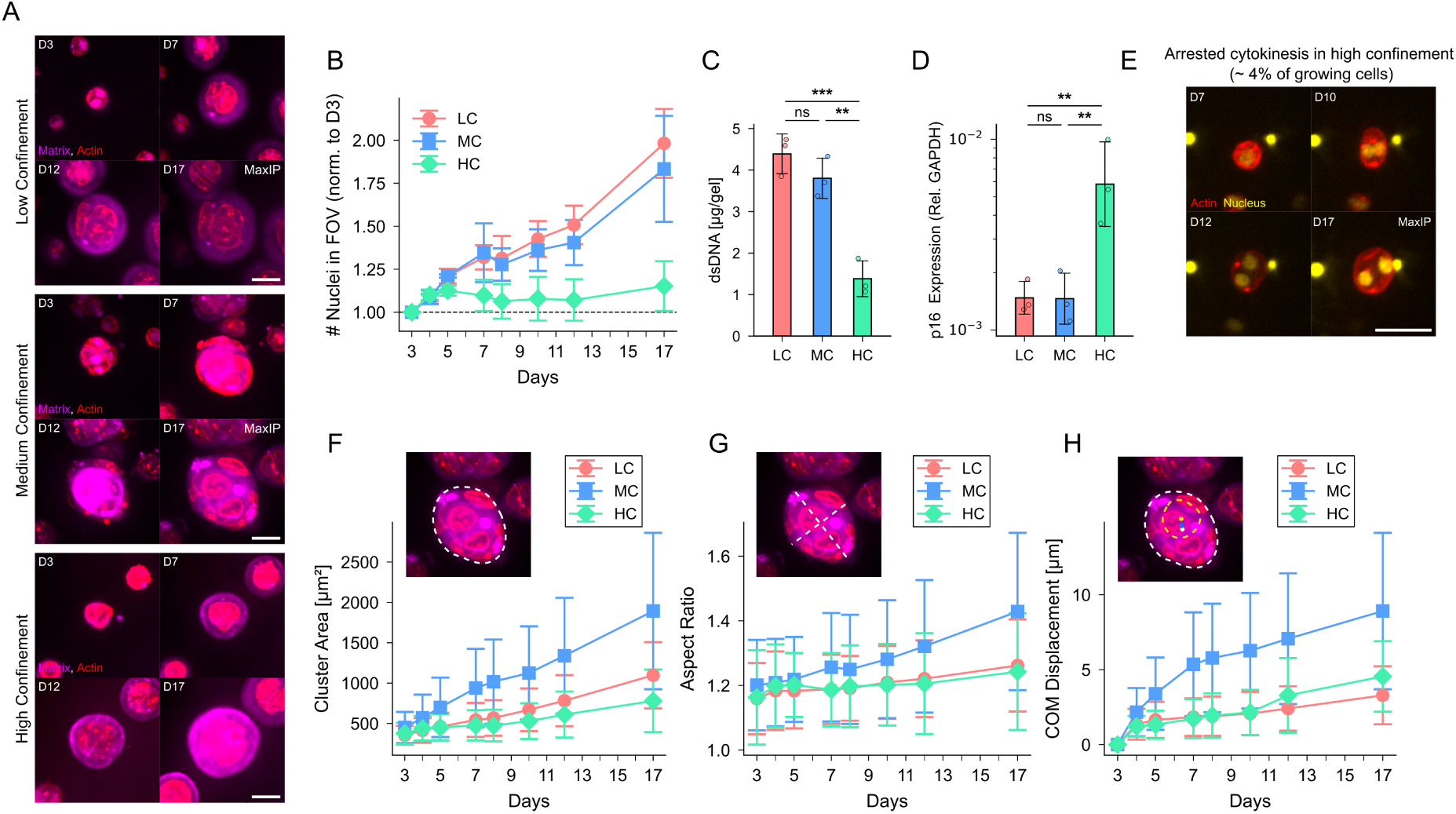
Effect of solid stress on chondrocyte cluster growth under medium and high confinement. **A)** Representative images of chondrocyte clusters growing in low, medium, and high confinement hydrogels. **B)** Proliferation quantified by nuclei count obtained from longitudinal time-lapse images, normalized to nuclei count at day 3. **C)** Mass of dsDNA in low, medium, and high confinement hydrogels at day 7. **D)** Relative gene expression of *p16* in chondrocytes in low, medium, and high confinement hydrogels. **E)** Representative time-lapse images of arrested cytokinesis in chondrocytes in high confinement hydrogels (see Fig. S10 for additional examples). **F, G, H)** Characterization of cluster area, aspect ratio, and center-of-mass displacement obtained from longitudinal time-lapse imaging. Scale bars 25 µm. N = 178, 203, and 37 for LC, MC, and HC for panels B, F, G, and H.

While chondrocytes in medium confinement hydrogels exhibited similar proliferation to those in low confinement hydrogels (Fig. 3B), the chondrocyte clusters in medium confinement hydrogels exhibited significantly different morphology compared to those in low confinement hydrogels. First, chondrocyte clusters in medium confinement hydrogels exhibited larger areas and higher aspect ratios compared to those in low confinement hydrogels (Fig. 3F, 3G). Furthermore, cluster shapes in medium confinement hydrogels also developed increased directionality during their growth, indicated by the displacement of the center-of-mass of the clusters in medium confinement hydrogels (Fig. 3H). These morphological features of clusters in medium confinement hydrogels can be attributed to higher cell-cell distance compared to those in low confinement hydrogels (Fig. 2J), rather than a difference in cell proliferation rates (Fig. 3B). While increased cell-cell distance could in theory arise from increased cell separation arising from nascent matrix secretions, our findings instead suggest the possibility of chondrocyte migration: cells are observed on the outside of the nascent matrix (i.e. at the cell-hydrogel interface) during the growth of clusters in medium confinement hydrogels (Fig. 3A), and indeed, intensity linescans of clusters reveal cells appearing to push out against the hydrogel, and occasionally leaving holes in the nascent matrix to do so (Fig. 1J, S11). Together, these findings suggest that medium confinement hydrogels impose a substantive level of solid stress onto the chondrocytes – enough to alter cell morphology and behavior, without resulting in cell cycle arrest and death – and promote directional growth of the cell clusters, potentially mediated by cell migration.

### Medium confinement induces mitosis-driven chondrocyte migration

We performed time-lapse imaging at shorter intervals to assess how directional cluster growth occurs in medium confinement hydrogels, and whether such migration occurs. While low confinement hydrogels facilitated an isotropic expansion and gradual proliferation of the chondrocyte clusters – showcasing a pivoting motion after cell division reminiscent of chondrocytes in the proliferative zone of cartilage^24^ – medium confinement hydrogels instead facilitated a striking migratory motion of the resident cells, in which the cells appeared to burst through the nascent matrix and push out towards the hydrogel wall (Fig. 4A, 4B, Movie 4, Movie 5). Approximately 73% of the observed clusters in medium confinement hydrogels exhibited these migratory behaviors (N = 282).

**Figure 4.**
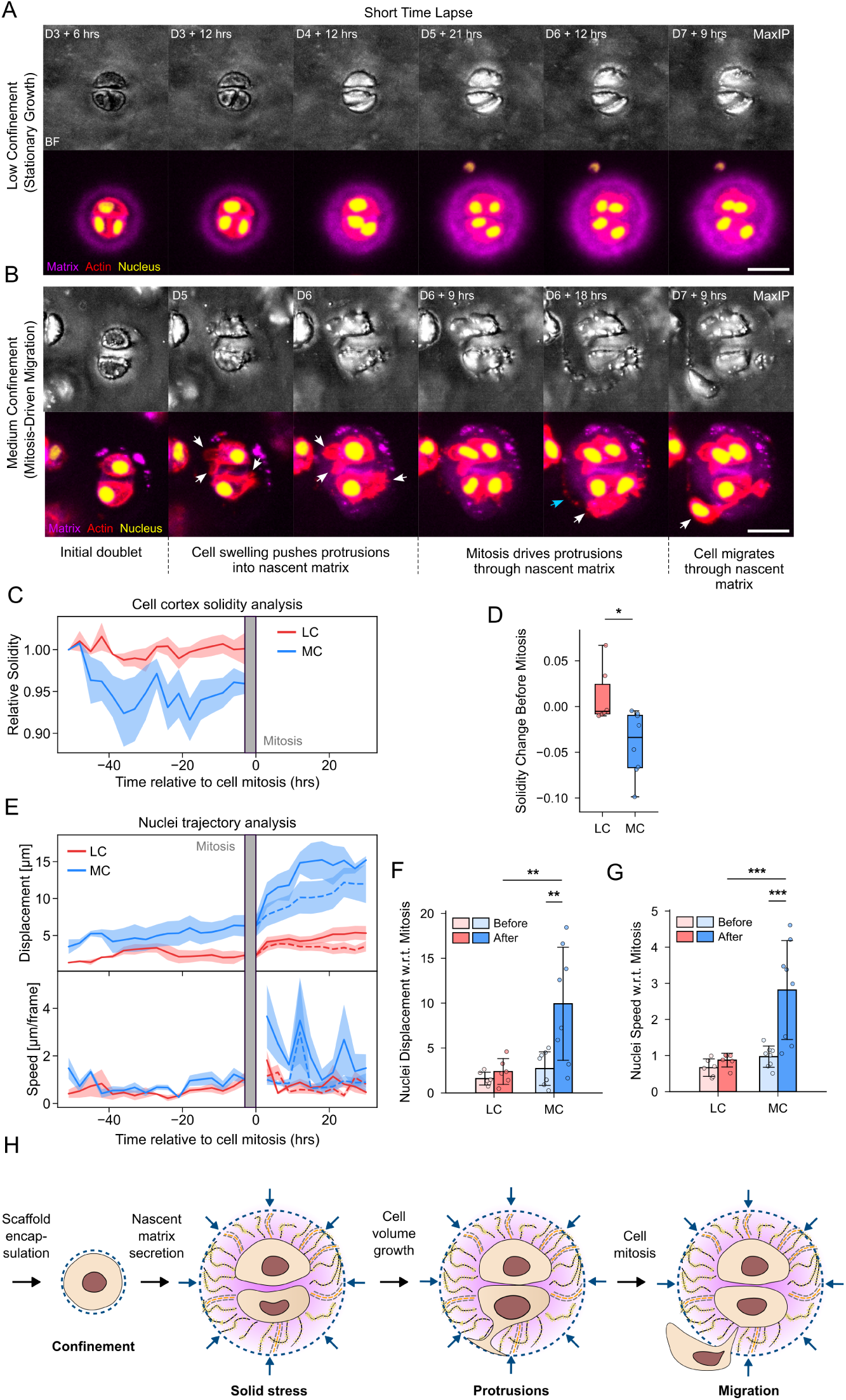
Solid stress mediates mitosis-driven chondrocyte migration through the nascent matrix. Representative time-lapse images of chondrocytes growing in **A)** low and **B)** medium confinement hydrogels. Protrusion expansion into the nascent matrix is highlighted by white arrows, and actin-rich strands are highlighted by blue arrows (see Fig. S13 for relevant discussion). Note that nascent matrix was labeled at day 3, and re-labeled at day 6. **C)** Relative change in morphological solidity in cells approaching cell mitosis. Solidity was normalized to the initial time point. **D)** Quantification of solidity change in cells approaching mitosis. **E)** Quantification of nuclei displacement and speed before and after mitosis. After mitosis, both of the daughter nuclei are tracked; solid and dashed lines indicate trajectories of the more and less motile nuclei, respectively. The nuclei displacement after mitosis is re-standardized to remove the contribution from mitosis itself. Each frame is 3 hrs. **F, G)** Average nuclei displacement and speed 20 hours before and after mitosis (more motile daughter only). **H)** Summary of the observed migratory dynamics of chondrocytes under medium confinement-induced solid stress. Scale bars 25 µm. N = 6 and 8 cells for LC and MC for panels C-G.

Mechanistically, this cell migration appeared to occur around mitosis. Specifically, we observed that cells extended protrusions into the nascent matrix during the volumetric growth phase, as evidenced by a ruffling of the cortex (or a decrease in cortex solidity) in cells approaching mitosis (Fig. 4C, 4D). Subsequently, mitosis appeared to push the protrusions as well as the daughter nuclei further outwards, ultimately promoting the migration of the daughter cell out of the nascent matrix and into the hydrogel (Fig. 4B); the resulting migration led to an increase in nuclei displacement and speed after cell mitosis (Fig. 4E, 4F, 4G). Altogether we found that cells under solid stress extended protrusions during volumetric growth, and used the outward force generated by mitotic cell division^23^ to migrate out of the nascent matrix (Fig. 4H). This migratory state of the cells in medium confinement hydrogels is independently evident from the cells migrating out of the nascent matrix upon hydrogel dissolution (Fig. 2B, Movie 1). Moreover, the migratory response appears to be independent of matrix degradation by metalloproteinases and aggrecanases, as similar protrusion-laden morphologies of cell clusters are observed even with pharmacological inhibition of matrix degradation (Fig. S12A, S12B).

Overall, chondrocyte migration under solid stress exhibits features that are reminiscent of amoeboidal modes of confined migration, such as adhesion-independent migration (into an alginate hydrogel that has no adhesive ligands), stress-driven growth of (bleb-like) protrusions into the matrix, and proteolysis-independent migration through the confining pores of the matrix.^25,26^ In this context, our results provide insight into how mitosis can propel amoeboid-like cells through their extracellular matrix. Additionally, during chondrocyte migration, we observe the nucleus squeezing through the narrow pores of the nascent matrix (Fig. S13A, Movie 6). This squeezing motion of the nucleus can create a piston-like effect that generates substantial pressures at the anterior of the cells, potentially resulting in the blebbing of the anterior membrane.^25,27,28^ Consistent with this idea, we observe strands of actin form at the hydrogel interface during nuclear migration through the matrix (Fig. 4B). Calcium imaging reveals that these strands are enriched in calcium, and hydrogel dissolution reveals that these strands indeed represent aggregates of small extracellular blebs (Fig. S13). Although the nuclear piston mode of migration is usually assumed to involve anterior matrix adhesion,^25,28^ we observe features of this mode in cells that migrate via mitosis into a hydrogel lacking adhesive ligands.

Lastly, the mitotic “bursting out” of the cells from the nascent matrix induces significant deformation of the hydrogel cavity, which was quantified by particle image velocimetry (PIV) analysis of the displacement of beads embedded in the hydrogels during the migratory events (Fig. S14, Movie 7). These cell-driven hydrogel deformations leave a persistent, crack-like feature in the cavity compared to the rounded cavity in low and high confinement hydrogels (Fig. S15). These findings show that the mitosis-driven migration of chondrocytes generates sufficient force to cause plastic yielding and fracture of the hydrogel cavity, which can allow cells to relieve the solid stress imposed on them from the nascent matrix confinement, and contributes to the directional growth of the chondrocyte clusters long-term (Fig. 3H). In summary, we observed a striking migratory behavior in chondrocytes under solid stress from nascent matrix confinement, where chondrocytes leverage mitosis to break through their nascent matrix and out towards the hydrogel (Fig. 4H).

### Medium confinement induces an inflammatory and osteoarthritic transcriptional profile

The observation of chondrocyte migration in Fig. 4 is surprising as chondrocytes are usually considered to be non-motile, as they are immobilized within the dense extracellular matrix of the cartilage. Yet, there have been emerging reports of migratory behavior both in normal juvenile chondrocytes^22^, and in chondroprogenitors (i.e., chondrocytes that exhibit stem-cell-like features) that emerge in cartilage tissue as a repair response to mechanical injury and osteoarthritis.^29–31^ To determine whether the migratory cells in medium confinement are associated with either physiological or pathological phenotypes, we studied gene expression related to cartilage homeostasis in cells from low and medium confinement hydrogels via RT-qPCR. We observed a general upregulation of catabolic markers such as *MMP3, MMP13, ADAMTS5* and *ADAM10*, as well as an upregulation in *COL2A1* – which encodes the main scaffolding protein of the articular cartilage – in cells under medium solid stress; this anabolic upregulation is also supported by quantification of total collagenous proteins secreted in the scaffolds via hydroxyproline assay (Fig. 5A). The simultaneous upregulation of catabolic and anabolic markers suggests that chondrocytes in medium confinement hydrogels exhibit a metabolically “active” state of remodeling their nascent matrix microenvironment. While some form of enhanced matrix remodeling can be observed during both postnatal growth of juvenile chondrocytes^22^ and early-stage osteoarthritis and injury especially in the context of progenitor activity,^30,32^ the specific upregulations of catabolic markers *MMP3* and *ADAMTS5* and inflammatory marker *IL6* are strongly associated with osteoarthritis and injury.^33–35^ These results provide initial evidence that cells under medium solid stress exhibit pathological phenotypes associated with osteoarthritis and injury.

**Figure 5.**
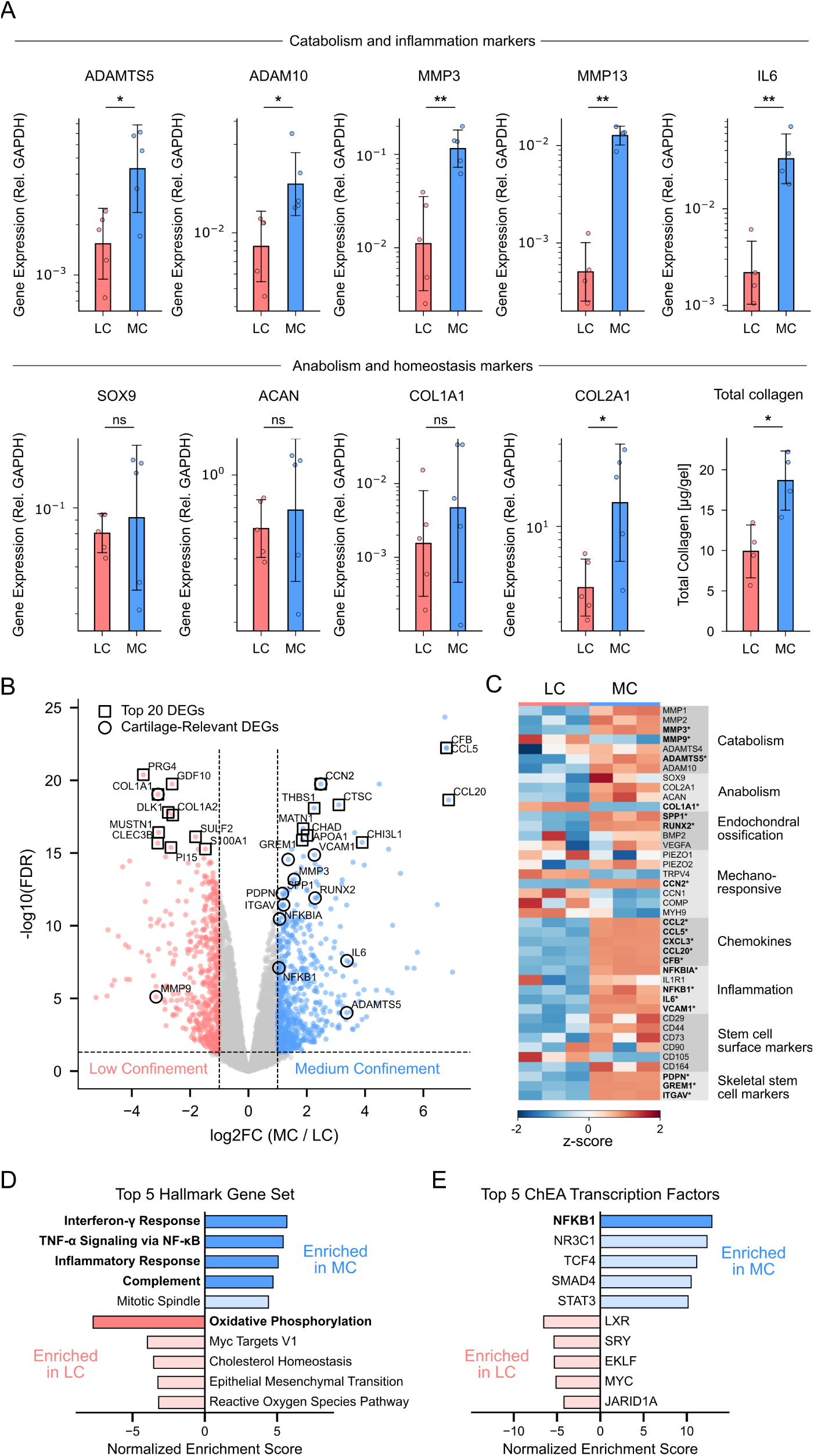
Injury- and osteoarthritis-associated transcriptional response of chondrocytes under solid stress. A) RT-qPCR gene expression of key catabolic, inflammatory, homeostatic, and anabolic genes relevant to chondrocytes, along with a total quantification of collagens in the scaffolds done via hydroxyproline assay, in low and medium confinement hydrogels. **B)** Volcano plot of differentially expressed genes (DEGs) (|log2 fold change| ≥ 1, adjusted p value < 0.05) between chondrocytes in low and medium confinement hydrogels. Top 20 DEGs and cartilage-specific DEGs are highlighted. **C)** Heatmap showing z-scores of selected cartilage-specific genes of relevance for individual repeats of LC and MC hydrogels. Genes in bold text are DEGs. A broad agreement between gene expression data obtained with RNA-seq and with RT-qPCR is observed (Fig. S16). **D)** Top 5 enriched Hallmark gene sets obtained via gene set enrichment analysis (GSEA) of the sequencing data. Bolded gene sets in medium confinement are associated with immune response and inflammation, and bolded gene set in low confinement relates to mitochondrial function. **E)** Top 5 enriched ChEA transcription factor gene sets. A significant enrichment of NFKB1, a master regulator of inflammation, is observed for medium confinement.

To examine the transcriptional response in greater detail, we performed RNA-seq on chondrocytes in low and medium confinement hydrogels. The results show that hallmark features of osteoarthritis and injury are indeed elevated in cells in medium confinement hydrogels, including simultaneous upregulation of catabolism and anabolism,^32^ upregulation of osteogenic markers,^36–39^ and elevated inflammatory signaling^40,41^ (Fig. 5B, 5C). The superficial zone protein marker *PRG4*, which serves a protective role against osteoarthritis initiation,^42^ is also significantly downregulated in cells in medium confinement hydrogels (Fig. 5B). Medium confinement hydrogels promoted a significant upregulation of chemokines such as *CCL2* and *CCL5*, and while we postulated that these chemokines could drive chondrocyte migration, the external addition of recombinant bovine CCL2 and CCL5 did not promote migratory dynamics in chondrocytes in low confinement hydrogels (Fig. S12C, S12D). Medium confinement hydrogels also promoted a general upregulation in MSC surface markers, as well as emerging skeletal stem cell markers such as *PDPN, GREM1, and ITGAV.*^43–45^ Notably, *GREM1* has also recently been implicated in mechanical-loading-induced chondrocyte inflammation via the *NF-κB* signaling pathway^46^. These results, alongside the migratory response observed in Fig. 4, suggest that solid stress on chondrocytes activates a progenitor-like state associated with osteoarthritis and injury.^29,31,47^

To further investigate the pathology associated with solid stress, we performed enrichment analyses of the transcriptional profiles of chondrocytes in low and medium confinement hydrogels. Enrichment analysis against the Hallmark gene sets showed that medium confinement hydrogels upregulated inflammatory and immune responses, and downregulated oxidative phosphorylation, both commonly observed features of osteoarthritis^40,48–50^ (Fig. 5D). These results were further reinforced by enrichment analysis against the ChEA transcription factor gene sets, showing a strong enrichment of the *NF-κB* pathway (regulator of inflammation) in medium confinement hydrogels (Fig. 5E). Altogether, transcriptional analyses implicate an upregulation of *NF-κB* inflammatory signaling in chondrocytes under solid stress arising from nascent matrix confinement, which accompanies chondroprogenitor-like phenotypic responses characterized by metabolic activation and migration. These results complement established understanding of *NF-κB* signaling as a hallmark response of compressive stresses,^51–53^ and suggest that solid stress can be a mediator of *NF-κB* signaling in chondrocytes. These findings have broad implications for understanding inflammatory responses in cartilage injury^46,54^ and osteoarthritis.^55,56^

## Discussion

In summary, we have demonstrated that the cell-secreted nascent matrix acts as a transducer of hydrogel mechanical confinement in 3D (Fig. 6). We showed that hydrogels can mechanically confine the cell-secreted nascent matrix at the cell-hydrogel interface, which imposes solid stresses onto the cells, and induces varying pathophysiological responses. While low solid stresses promoted a proliferative state and high solid stresses induced cell cycle arrest and death, medium solid stresses induced a chondroprogenitor-like response characterized by co-elevation of anabolism and catabolism, inflammatory signaling with enriched *NF-κB* transcriptional targets, upregulation of stem cell markers, and a striking migratory response centered around mitosis. We also showed that this confinement-induced solid stress can be tuned by remodelable properties of the hydrogel such as viscoelasticity and plasticity, as hydrogel remodeling facilitates penetration of the secreted nascent matrix into the hydrogel, rather than confinement at the cell-hydrogel interface. These results provide new insight into chondrocyte pathology under solid stress, and demonstrate tunability of these responses *in vitro* through hydrogel materials design.

**Figure 6.**
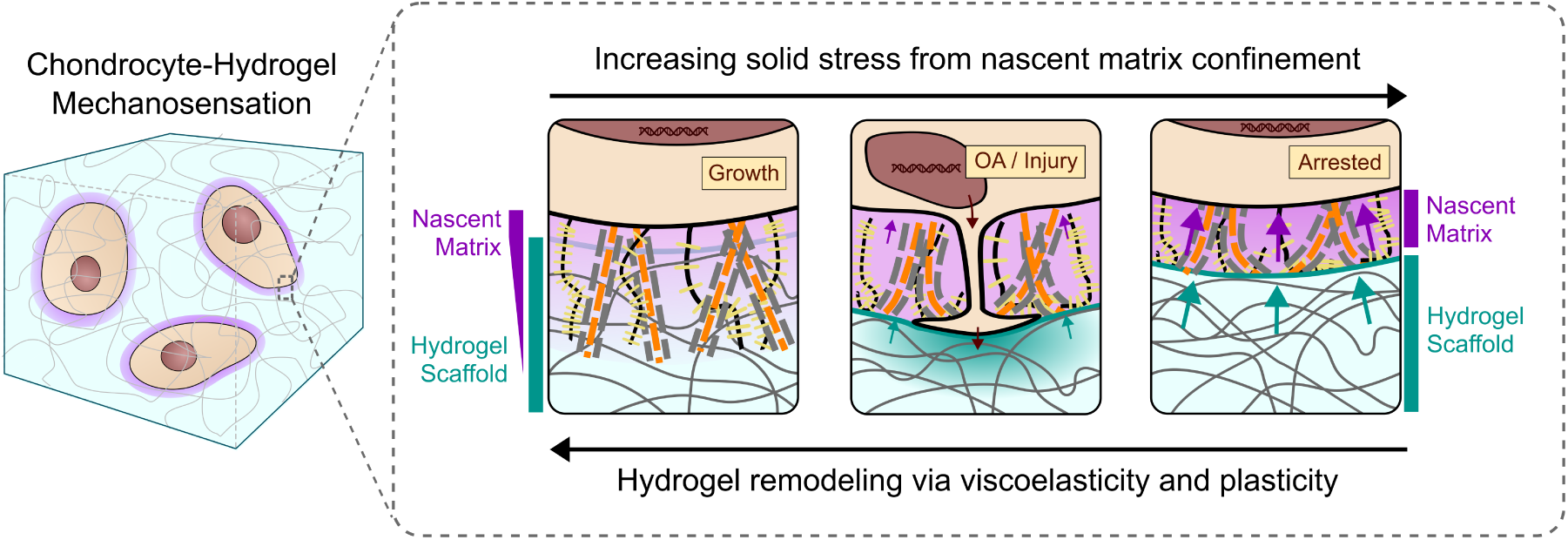
Hydrogel mechanical properties regulate nascent matrix confinement, solid stress, and pathophysiology of chondrocytes. High hydrogel remodelability (via viscoelasticity and plasticity) facilitates penetration of the nascent matrix into the scaffold, whereas low hydrogel remodelability instead confines the nascent matrix at the scaffold interface, imposing solid stress on resident chondrocytes. Low solid stresses facilitate a homeostatic and proliferative response. Medium solid stresses induce phenotypes associated with cartilage osteoarthritis and injury, including inflammation via enrichment of *NF-κB* signaling pathway, activation of both anabolic and catabolic programs (Fig. 5), and a chondroprogenitor-like migratory response where cells leverage mitosis to breach their nascent matrix (Fig. 4H). High solid stresses cause cell cycle arrest and death in chondrocytes.

These findings provide important context to prior studies on nascent matrix secretions in hydrogel scaffolds. Previous studies showed that while the nascent matrix provides important biological ligand signals for the spreading and differentiation of MSCs, both nascent matrix assembly and cell fate remained sensitive to the mechanical cues of the confining hydrogels such as elasticity, viscoelasticity, and degradability.^9,11^ Increasing the stiffness of the hydrogels was associated with a more compacted layer of nascent matrix, an effect which was also correlated to nascent matrix remodeling and altered cell differentiation.^9,10^ Our results suggest these effects on cells may be downstream of the solid stress that is applied by nascent matrix confinement, which is mediated by the mechanical properties of the hydrogels such as stiffness, viscoelasticity, and plasticity. Similarly, our findings may also explain the more diffuse and penetrated structure of the nascent matrix secreted by chondrocytes and MSCs in agarose hydrogels^57^ – which exhibit fast viscoelastic stress relaxation similar to our low confinement case where matrix penetration was observed^4^ – compared to those observed in covalently cross-linked hyaluronic acid hydrogels.^10^ Lastly, our results provide biophysical context for the observation of protrusions in chondrogenic MSCs in hydrogel scaffolds, as a consequence of competition between hydrogel-mediated solid stress and volumetric growth of cells.^58^ Altogether, our results demonstrate that hydrogel confinement remains a critical variable for directing the fate of matrix-secreting cells, through nascent matrix confinement and subsequent exertion of solid stress on cells. These hydrogel-matrix interactions should therefore be considered a fundamental design parameter for biomaterials used in tissue engineering and disease modeling applications. As such, while our work establishes the role of hydrogel mechanics in mediating this interaction, establishing how other material properties such as proteolytic degradability, chemical functionality, and porosity may alter this interaction represents an important area of future research for biomaterials science.

The demonstration that scaffold-mediated solid stresses promote a *NF-κB* transcriptional response in chondrocytes, alongside inflammatory and progenitor-like phenotypes observed during osteoarthritis and injury, may be relevant to the *NF-κB* signaling that has been reported in cartilage injury^46,54^ and osteoarthritis.^55,56^ In cartilage injury caused by excessive mechanical loading, the cells experience direct compressive stresses from the loaded solid matrix.^59^ In chronic osteoarthritis, it is known that the matrix undergoes cross-linking via the accumulation of biomarkers such as advanced glycation end-products (AGEs)^60,61^ and lysyl oxidase (LOX),^62^ which results in increased stiffness, reduced viscoelastic dissipation,^63,64^ and reduced susceptibility to proteolytic degradation^65^ – all of which can lead to an increase in solid stress on the cells. Taken together, solid stress – a concept normally used to describe the compressive stress state of tumors^17,66^ – could also serve as a relevant framework for describing chondrocyte mechanobiology in the context of osteoarthritis and injury. Relatedly, while we have demonstrated that chondrocytes exhibit a progenitor-like response under solid stress, including upregulation of stem cell markers as well as a migratory response, an important follow-up study would be to assess whether these cells also exhibit multilineage capability. These validations would be an important step towards establishing a robust method for generating and studying chondroprogenitors *in vitro* via hydrogel confinement.

Lastly, these findings highlight a new mitosis-driven mechanism of confined migration for cells under matrix confinement and solid stress. Although the migratory ability of chondrocytes and chondroprogenitors has been known, and even observed in alginate cultures in the past,^67,68^ it has remained unclear as to how these cells could actually migrate under confinement.^69^ Our work provides a plausible biophysical mechanism by which such chondrocytes might be able to undergo migration in the presence of matrix confinement, via stress-induced protrusion formation and mitosis-driven migration. This mechanism may also be relevant for the confined migration of other cells which are matrix-secreting, proliferating, and under solid stress – for instance cancer cells in tumors^66^ – and in this sense, our findings emphasize the importance of studying the confined migration of cells in 3D environments where such behaviors can be accessible. Although our work provides initial insight into how chondrocytes leverage known mechanisms of confined migration during their mitosis-driven migration – including features associated with amoeboidal migration and the nuclear piston – the exact biophysical machinery that underlies the solid stress-driven protrusion formation and mitotic migration across different tissue contexts, and the cell biological consequences of such migratory decisions, remain important future questions for broadly understanding confined migration in tissues.

## Acknowledgement

J.S. acknowledges financial support from the NIH F32 Postdoctoral Fellowship via F32AR084286. N.B. and O.C. acknowledge financial support from R01AR081993.

## Methods

### Reagent Source

All reagents and appropriate abbreviations, as well as their sources, are listed in Table S1.

### Chondrocyte Isolation

Chondrocyte isolation was performed following a slight variation of the protocol in ref.^4^ Hyaline articular cartilages were harvested from the femoral groove of the legs of juvenile (1-3 weeks old) bovine calves acquired from commercial suppliers (Research 87 and San Jose Valley Veal). The cartilage on the femoral groove was rinsed in DPBS containing 1 X Gentamicin-Amphotericin and the topmost hyaline layer of cartilage was harvested (in order to avoid subchondral bone, and regions with increased vasculature). The hyaline layer was thinly sliced and diced into small ∼ 1 mm^3^ cubes, after which they were placed in a digestion medium containing DMEM, 1 mg/mL collagenase 2, 1 mg/mL collagenase 4, and 1 X Anti-Anti and placed in a tissue culture incubator. The digestion medium was gently agitated over 48 hours. After digestion, the digestion medium and cartilage tissues were filtered with a 40 µm cell strainer, and then centrifuged (500 g, 7 min), aspirated, and resuspended in DPBS twice. The final cell count was determined using a VI-CELL (Beckman Coulter), and then frozen in 10 million cells / mL aliquots, containing 50 % FBS, 10 % DMSO, and 40 % cell culture media (see Cell Culture).

### Fluorescent Alginate Synthesis

Fluorescent alginate synthesis was performed following a slight variation of the protocol in ref.^71^ Low or high molecular weight alginates were first dissolved in MES buffer at pH 6.5 at 1 wt. % concentration, after which Sulfo-NHS and EDC were sequentially dissolved in the mixture at 10 % molar equivalent of the uronic acid in the alginate solution. Then, 6-aminofluorescein was dissolved in 1 mL of dimethyl sulfoxide (DMSO), and added to the reaction mixture at a final concentration of 1.25 mM per 1 wt. % of alginate (stoichiometric ratio of 2 fluorescein groups per polymer chain). All calculations assumed 60% coupling efficiency. The reaction was left to proceed for 24 hours, after which it was quenched with hydroxylamine hydrochloride, dialyzed in a 3500 MWCO dialysis membrane against DI water initially containing 150 mM NaCl; the NaCl concentration was gradually reduced to 0 mM across 10 dialysate exchanges over 3 days. Afterwards, the dialyzed solution was treated with activated charcoal, sterile-filtered, and lyophilized over one week.

### Cell Encapsulation in Hydrogels

Prior to encapsulation, all chondrocytes were thawed, rinsed in DPBS (i.e., re-suspension, centrifugation (500 g, 7 min), and supernatant aspiration), re-suspended in cell culture medium, and plated on tissue culture flasks overnight. On the next day, cells were released using trypsin-EDTA for 5 minutes in the incubator, after which the suspension was rinsed first in a solution containing 50% cell culture media and 50% DMEM, filtered using a 40 µm cell strainer, and then rinsed again in DMEM. Then, the cells were counted using VI-CELL and then re-suspended in a small volume of DMEM at a concentration of approximately 30 million cells / mL.

Alginate hydrogels (low and medium confinement hydrogels) were made following protocols in ref^12^. This was done via syringe-mixing, where a Luer lock coupler was used to mix one syringe containing alginate-DMEM solutions with cell suspensions with another syringe containing DMEM and CaSO_4_. The mixture was deposited on a glass plate that was hydrophobically treated in advance with Sigmacote, and covered with another glass plate with 2 mm spacers between the plates. The hydrogel was allowed to gel for 45 minutes, after which the gels were placed in media and then in the incubator. For regular culture, gels were made into 5 mm discs using a biopsy punch, and each disc was placed in a well of a 24-well plate. For time-lapse imaging experiments, gels were directly cast within 24-well glass-bottom tissue culture plates.

Poly(ethylene glycol) (PEG) hydrogels (high confinement hydrogels) were made following variations of the protocols in ref.^13^ A precursor mixture containing 8-PEG-norbornene, 2-PEG-thiol, LAP, and the cell suspension was either sandwiched between glass plates or directly cast within 24-well glass-bottom tissue culture plates, in similar vein as the alginate hydrogels. The precursor was then cured with a standard commercial 405 nm LED resin-curing lamp for 3 mins. We note that live-dead experiments using 1:1000 Calcein-AM and ethidium homodimer-1 were performed after light curing to verify that the curing process did not lead to adverse cell viability outcomes. After curing, gels were placed in media and then in the incubator.

All hydrogel encapsulations were done at a 3 million / mL cell density. Low confinement hydrogels consisted of 2 wt. % low molecular weight alginate and 28 mM CaSO_4_, and medium confinement hydrogels consisted of 2 wt. % high molecular weight alginate and 10 mM CaSO_4_. High confinement hydrogels consisted of a total of 4 wt. % 8-PEG-norbornene and 2-PEG-thiol at a 1:3 stoichiometric ratio of 8-PEG-norbornene to 2-PEG-thiol, and 0.05% w/v of the photoinitiator LAP. Fluorescent variations of the low and medium confinement hydrogels were made by simply substituting regular alginates with the synthesized fluorescent alginate. Fluorescent variations of the high confinement hydrogels were made by adding a small amount of FITC-PEG-thiol to the mixture, at a 1:25 stoichiometric ratio of FITC-PEG-thiol to 2-PEG-thiol. When preparing fluorescent hydrogels, all curing processes are done under protection from ambient light with foil.

### Cell Culture

Chondrocytes were cultured following established protocol.^4^ Hydrogel-encapsulated cells were cultured in medium consisting of low-glucose DMEM, 10 % FBS, 0.1 mM MEM-NEAA, 10 mM HEPES, 0.05 mg / mL L-ascorbic acid-2-phosphate, and 1 mM CaCl_2_. The medium was changed every 2-3 days.

Chondrocyte culture experiments involving metabolic labeling of the nascent matrix were done following labeling protocols in refs.^10,72^ Separate cell culture media were created for labeling nascent proteins (with L-azidohomoalanine, AHA) and nascent glycans (with Ac4ManNAz, MNZ); we note that we have also tried labeling glycans by substituting a different sugar group, Ac4GalNAz,^16^ and observed similar labeling results as those with Ac4ManNAz.

For MNZ medium, 10 ∼ 50 µM of MNZ were added to the regular cell culture medium (we initially started at 10 µM, but found no adverse effects at 50 µM). For AHA medium, the low-glucose DMEM was substituted to a high-glucose DMEM without glutamine, methionine, cystine, and sodium pyruvate. To this medium, L-glutamine, L-cystine, sodium pyruvate, and 100 µM L-methionine were individually re-added to match the original concentrations in DMEM. As the cystine is dissolved in 1M HCl,^72^ an equivalent amount of 1M NaOH was added to balance the pH. Lastly, 100 µM of AHA was added. All media changes were done with either AHA or MNZ medium, depending on the labeled nascent matrix of interest.

### Rheology

All mechanical characterizations were performed using a TA DHR-3.

Storage modulus *G*′ and tan *δ* were characterized on hydrogel discs which were 8 mm in diameter and 1 mm in thickness, and which were gelled for 1 hr and equilibrated in cell culture medium for 24 hours overnight. The gels were loaded on a rheometer, and a 8 mm parallel plate was brought down onto the gel until a positive normal force of 0.02 N was registered. The gel was allowed to equilibrate for 5 minutes before small amplitude oscillatory shear experiments at an angular frequency of ω = 1 rad/s were performed for 3 minutes, and the final *G*′ and tan *δ* values are reported in Fig. 1C and 1D. All experiments were done at room temperature.

Stress relaxation experiments were performed by curing the hydrogels *in situ* on the rheometer plate under a 20 mm parallel plate geometry, with the samples covered in mineral oil. Gelation was allowed to proceed for 1 hr at room temperature. Stress relaxation experiments were then performed by imposing a 1 % shear strain for up to 5000 s. The obtained relaxation moduli *G*(*t*) were normalized to a reference modulus at *t* ∼ 4 s to allow for strain equilibration, as illustrated in Fig. 1E. Afterwards, the data were fitted to a Kohlrausch-Williams-Watts stretched exponential function, *G*(*t*)/*G*(*t_ref_*) = exp{−(*t*⁄*τ*)*^β^*}, where *τ* is the characteristic relaxation time and *β* is the stretching exponent. *β* was fixed to 0.3, which is a common exponent found in the stress relaxation of many soft materials^73^, and was found to fit our data well (Fig. S1). The stretched exponential function implies a distribution of relaxation modes, and so we computed a mean relaxation time 〈*τ*〉 = (*τ*⁄*β*) Γ(1⁄*β*) where Γ is the Gamma function.^74^ The inverse of this term, the mean relaxation rate 〈*k*〉, is reported in Fig. 1F.

*In situ* experiments on high confinement hydrogels required a special setup, as gelation required 405 nm light irradiation. To facilitate this, a custom 3D printed box consisting of a non-UV-absorbing glass top plate and the 405 nm lamp housed inside was used so that irradiation could be performed from below the sample. The 3D printed box was secured onto the rheometer by using adhesive-backed sandpapers (Trizact A10) on the box as well as the rheometer bottom plate, and bonding these sandpapers together using cyanoacrylate superglue, which ensured a firm hold. Gelation was allowed to proceed under light irradiation for 3 minutes, while the sample was sealed with mineral oil, after which the sample was allowed to equilibrate for 1 hr. Stress relaxation experiments were then performed by imposing a 1 % shear strain for 1000 s; longer experiments were not performed as no relaxation was observed.

Amplitude sweep experiments were performed at ω = 1 rad/s also on *in situ* gelled samples. High confinement hydrogels required the use of sandpapers on the geometry as well as the glass bottom to prevent slip (the semi-transparency of the Trizact A10 sandpaper allowed light irradiation to occur through the sandpaper). The yield strain and stress were obtained from deviations in the loss modulus *G*′′ from the baseline, as discussed in detail in Fig. S2.

### Labeling and Fluorescent Imaging

Fluorescent imaging experiments were performed using two confocal microscopes – a Leica SP8 confocal microscope and a Nikon Ti2-E confocal microscope, both loaded with temperature, humidity, and CO_2_ controlled environments for live imaging. A 20x 0.75-NA dry objective was primarily used for the Nikon (occasionally, a 10x 0.45-NA dry objective was also used, and noted as such on the relevant images), and a 20x 0.75-NA dry objective or 25x 0.95-NA wet objective was used for the Leica.

For labeling of the cell cortical actin, a SiR-Actin fluorescent probe was added to the cell culture medium at a 1:10000 dilution during live imaging. For labeling of the cell nuclei, a SPY-555 fluorescent probe was added to the cell culture medium at a 1:5000 dilution during live imaging. Both of these probes are specific and do not need to be washed out after labeling.

For labeling the nascent matrix, we found that the relevant fluorescent probe that binds to the azide-modified proteins or glycans, DBCO-555, exhibited non-specific binding unless the probe was rinsed out. We thus followed a variation of the established protocol for labeling the nascent matrix,^10,11,72^ with minor adjustments. The hydrogels were rinsed in Tris-buffered saline with 3 mM of CaCl_2_ (cTBS) (where the calcium is added to match equivalent concentrations in DMEM and prevent alginate gel dissolution) and labeled with a 30 µM solution of DBCO-555 in cTBS buffer containing 1% BSA for 1 hr. After staining, the gels were thoroughly rinsed with a cTBS-BSA solution once (15 minute equilibration), DMEM once (15 minute equilibration), and then cell culture medium twice (1 hour equilibrations each). This labeling process was performed every 2-3 days during time-lapse imaging.

A typical confocal imaging experiment collected z-stack images of the fluorescent hydrogel (488 nm laser), the nascent matrix (555 nm laser), the cell (647 nm laser), and the brightfield. In some cases where we imaged both the cell nuclei and the nascent matrix, it is noted that SPY-555 and DBCO-555 shared a laser line. However, the labeling differences were conspicuous, with bright nuclei-shaped areas stained within cells for SPY-555 and a diffuse signal outside of cells for DBCO-555. Thus, the nuclei were segmented using the nuclei pretrained model in Cellpose (manually reviewed for accuracy), and labeled with different colors in the figures.

### Intensity Linescans

All linescans were done by drawing a small, elongated rectangle across cross-sectional images of cell clusters to compute the spatially averaged intensity linescans of the different channels on ImageJ. First, the hydrogel intensity profile was smoothed, and the outer regions were used to estimate baseline intensity. The left and right cavity walls were defined to be at half-depth thresholds between this outer baseline and the minimum intensity within the dip. Each linescan was split at the center of the cavity, and both sides were aligned to their corresponding wall and interpolated onto a common distance grid spanning -20 µm (cavity side) to +20 µm (hydrogel side) of the wall. All channel intensities were then scaled using channel-specific reference values. For the hydrogel channel, the cavity-side baseline was set to 0 and the outer baseline was set to 1. For the matrix and cell channels, the outer baseline was set to 0 and the mean intensity within the cavity was set to 1. Individual linescans of different clusters are averaged and shown as mean ± standard error of mean (SEM) in Fig. 1J.

### Image Segmentation and Analysis

Segmentation of cells and nuclei in cross-sectional images was done by using cyto3 and nuclei pretrained models in Cellpose. Segmentation of cells and the nascent matrix from 3D z-stacks was performed on Imaris, and associated morphological metrics such as oblate ellipticity and volume were also obtained via Imaris. Segmentation of whole cell clusters were manually performed using the Cellpose GUI. All segmentations were manually reviewed for accuracy, and were subsequently used for morphological analysis.

All time-lapse images in this work are pre-processed by performing StackReg with respect to either the cell or brightfield channel to correct for the drift, and then performing bleach correction to correct for photobleaching effects.

### Dissolution Assays

All dissolution assays were performed by first placing discs of cell-laden hydrogels in wells of Lab-Tek 8 well slides (Nunc) and curing a low-gelling temperature agarose around and over the discs to prevent hydrogel displacement and drift during perturbation experiments. For hydrogel dissolution experiments, either alginate lyase in DMEM (1.5 mg / mL) or EDTA dissolved in DPBS at a pH of 7.4 (50 mM) were added to the Lab-Tek wells while performing live imaging. For matrix dissolution experiments, collagenase 4 in DMEM (1.5 mg / mL) was added. We note that, despite the nomenclature “collagenase”, collagenase enzymes can indiscriminately degrade nascent matrix components, including proteoglycans. Confocal z stacks were obtained with time to ensure that the same cluster was imaged during the dissolution experiments.

### Particle Image Velocimetry

For particle image velocimetry (PIV) analysis of bead displacements during chondrocyte migration in medium confinement, 0.2 µm carboxylated fluorescent microparticle solutions are added to the hydrogel precursor mixture before gelation (2.5 µL / mL of the 2 % w/v stock). Time-lapse cross-sectional images at day 7 were first pre-processed with StackReg and bleach correction, and then PIVlab (http://PIVlab.de) was used in MATLAB to track bead displacement. Cumulative displacement fields were computed by direct correlation between the first frame and each subsequent frame.

### Qubit Assay

To extract DNA from hydrogels, each 5 mm diameter 2 mm thick cell-laden hydrogel was disintegrated using an Eppendorf-compatible micropestle in a 100 µL solution of genomic lysis buffer, and frozen in - 20 °C and stored. For quantifying the amount of dsDNA in the hydrogels, 10 µL aliquots of these lysis solutions were analyzed with a Qubit 1x dsDNA broad range assay kit according to manufacturer protocol. Sample fluorescence at 485/530 nm was measured by using a Synergy H1 microplate reader (Agilent) to quantify dsDNA. The fluorescence response from the hydrogels was normalized to the baseline response of 10 µL of the genomic lysis buffer. All dsDNA readings were scaled back to the 100 µL lysate volume to obtain a per-gel reading. All analysis was done in triplicates.

### Hydroxyproline Assay

The total content of collagen in hydrogels was calculated via hydroxyproline assay, following established protocol.^4^ First, hydrogels were digested in DPBS containing papain at 60 °C overnight. Subsequently, the papain digested supernatants were used to quantify hydroxyproline content. Briefly, this involves performing hydrolysis in concentrated HCl, desiccating the samples, performing oxidation with a chloramine-T oxidation buffer, and then adding Ehrlich’s aldehyde containing DMAB to spectrophotometrically detect hydroxyproline. Hydroxyproline was converted to collagen amount with a 1:7.46 hydroxyproline:collagen mass ratio.

### Reverse-Transcription Quantitative Polymerase Chain Reaction

To extract RNA from hydrogels, two 5 mm diameter 2 mm thick cell-laden hydrogels were disintegrated using an Eppendorf-compatible micropestle in 300 µL of RNA lysis buffer, and frozen in -70 °C and stored. At least three of these vials were collected to ensure all analysis was done in triplicates or more. RNA was then extracted using the Quick-RNA Microprep kit, following manufacturer protocol. Next, reverse transcription was carried out to generate complementary DNA using PrimeScript RT Reagent Kit (Takara Bio). Reverse-transcription quantitative polymerase chain reaction (RT-qPCR) was performed by using a Power SYBR Green Master Mix. All gene expression values were normalized against the GAPDH housekeeping gene (ΔCt). All primers sequences (IDT) are listed in Table S2.

### Bulk RNA-Sequencing

All RNA-seq experiments on low and medium confinement hydrogels were done by first releasing the cells from the alginate hydrogels with EDTA dissolved in DPBS at a pH of 7.4 (50 mM), after which the EDTA was rinsed, and then the cells were pelleted. Cell pellets were stored in DNA/RNA Shield, and triplicates of low and medium confinement conditions were submitted to Plasmidsaurus for RNA-sequencing. Differentially expressed genes (DEGs) were identified from vendor-supplied edgeR results, and defined as genes with |log2 fold change| ≥ 1, and adjusted p value < 0.05. Gene set enrichment analysis (GSEA) was performed via blitzGSEA on genes pre-ranked by log2 fold change against the MSigDB Hallmark 2020 library and the 2022 ChEA transcription factor gene library.

### Statistics

All statistical data for the study are collected across at least three hydrogel encapsulation replicates. All statistical analyses were performed using Python (SciPy). Statistical significance between two groups was done with a Welch’s t-test, and between three groups or more was done with a one-way ANOVA with Tukey’s honestly significant difference post-hoc test. * p < 0.05; ** p < 0.01; *** p < 0.001. Shaded error bars denote standard deviations, except in two instances – linescan intensities in Fig. 1J and migration analysis in Fig. 4C and 4E, where shaded error bars denote standard error of mean, for the sake of data clarity.

