## Supplementary Information for "Hydrogel confinement of the secreted nascent matrix imposes solid stress and mediates pathology of chondrocytes"

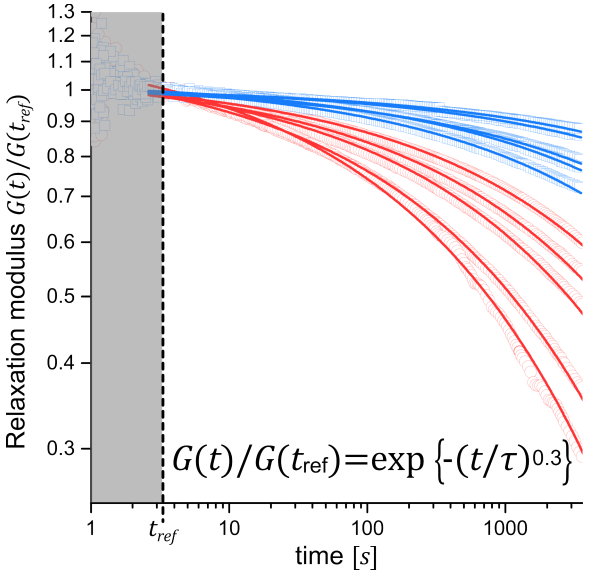

**Figure S1. Kohlrausch-Williams-Watts (KWW) fit of the stress relaxation data.** Kohlrausch-Williams-Watts stretched exponential function exhibits a functional form $G(t)/G(t_{ref}) = \exp\{-(t/\tau)^{\beta}\}$, where $\tau$ is the characteristic relaxation time and $\beta$ is the stretching exponent. $\beta$ was fixed to 0.3, which is a common exponent found in the stress relaxation of soft materials[1], and was found to fit our data well. The stretched exponential function implies a distribution of relaxation modes, and so we computed a mean relaxation time $\langle\tau\rangle = (\tau/\beta)\,\Gamma(1/\beta)$ where $\Gamma$ is the Gamma function.[2] The inverse of this term, the mean relaxation rate $\langle k\rangle$, is reported in Fig. 1F.

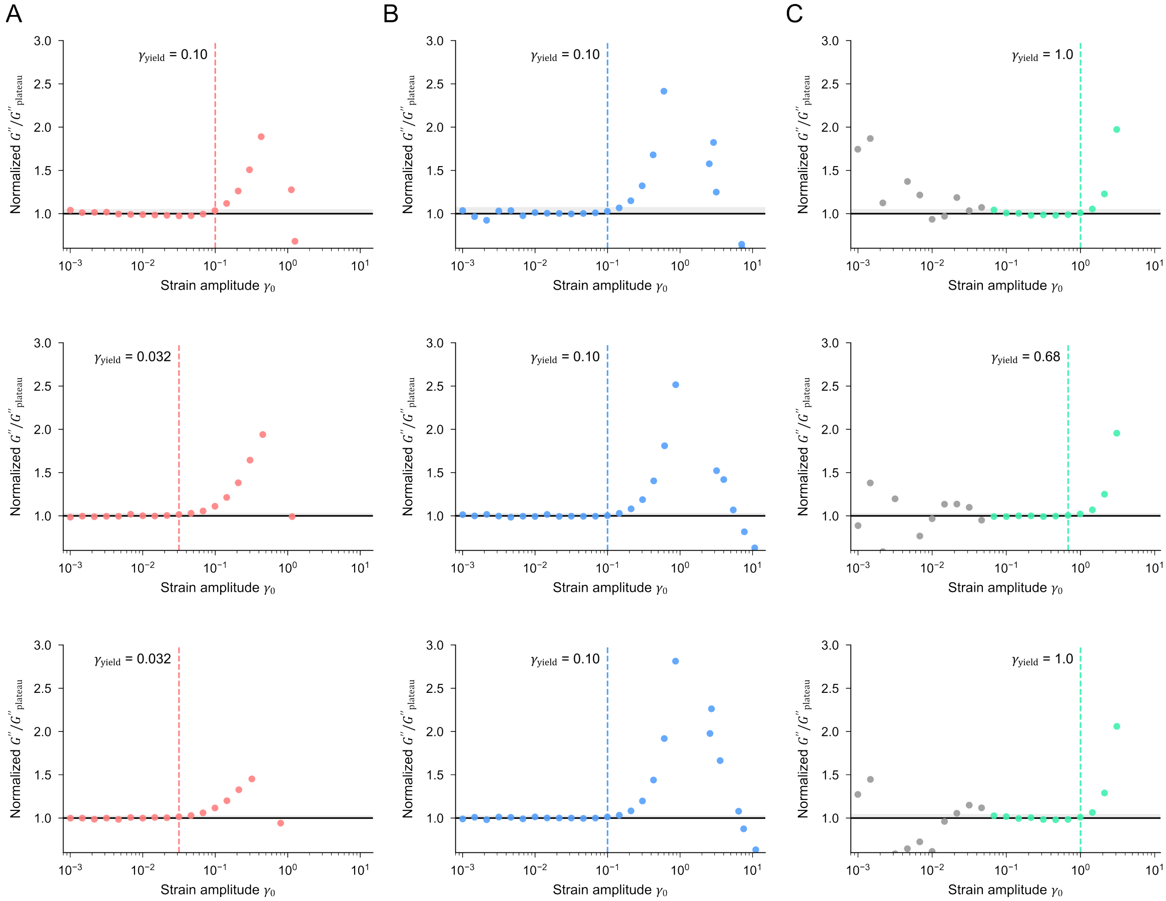

**Figure S2. Protocol for quantifying yield strain from increases in the loss modulus.** Loss modulus $G''$ normalized to the average of the plateau for A) low confinement, B) medium confinement, and C) high confinement hydrogels. The yield strain $\gamma_y$ is defined as the applied strain amplitude $\gamma_0$ beyond which the normalized $G''$ exceeds two standard deviations of the plateau (marked as a shaded grey band). The increase in $G''$ is a hallmark signature of the accrual of unrecoverable plastic deformations in the material under large amplitude oscillatory deformations.[3] The noisy data at low $\gamma_0$ for high confinement gels (marked as grey symbols) are excluded from the plateau.

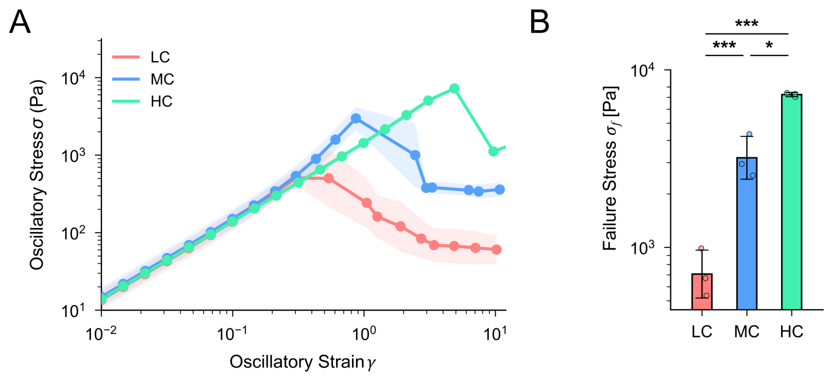

**Figure S3. Characterization of ultimate failure stress from amplitude sweeps.** A) Averaged traces of the raw oscillatory stress $\sigma$ as a function of raw oscillatory strain $\gamma$ during amplitude sweep experiments. The maximum oscillatory stress corresponds to the point of macroscopic failure in the gel, which was also verified by the visible breakage of the gel upon unloading the gel from the rheometer. N = 3 for all three conditions. B) The accordingly defined failure stress $\sigma_f$ in low, medium, and high confinement hydrogels.

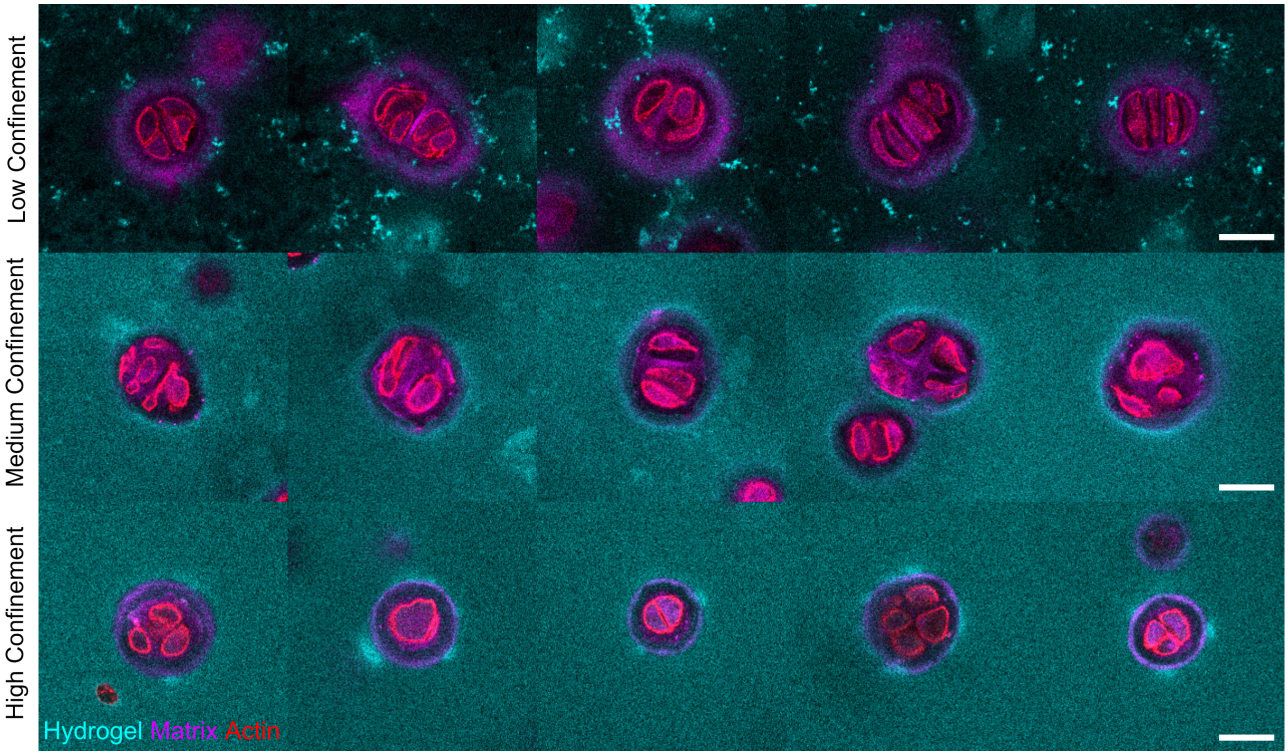

**Figure S4. Additional images of multicellular chondrocyte clusters at day 7 across low, medium, and high confinement hydrogels.** Five representative examples are shown per condition. Scale bars 25 µm.

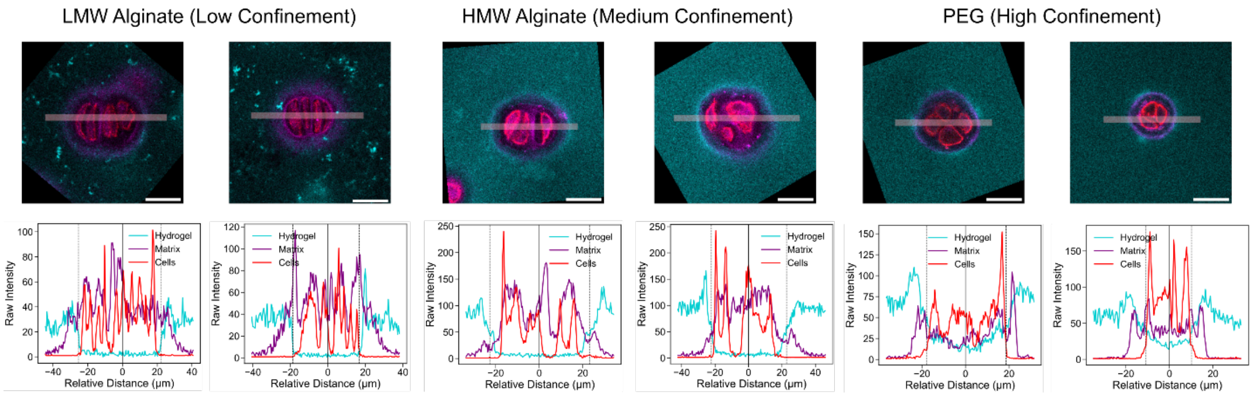

**Figure S5. Example raw traces of intensity linescans of chondrocyte clusters.** Images are obtained at day 7 across low, medium, and high confinement hydrogels. Note that for Fig. 1J, the left and right side scans of these raw intensity traces are folded onto one common axis that quantifies the relative distance from the hydrogel wall. Two representative examples are shown per condition. Scale bars 25 μm.

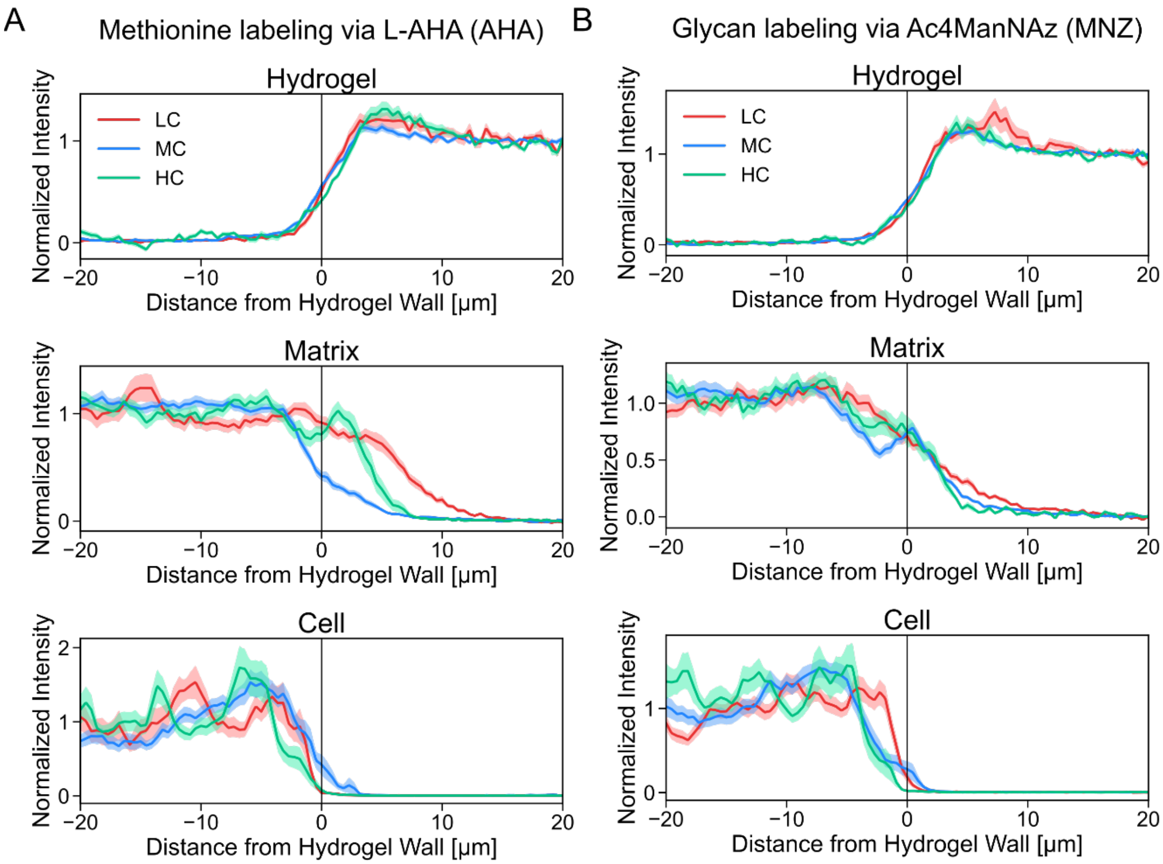

**Figure S6. Comparison of intensity linescans between nascent proteins and nascent glycans.** Nascent proteins are labelled with AHA, and nascent glycans are labelled with MNZ. The penetration of MNZ-containing nascent matrix is less distinguishable compared to AHA. N(AHA) = 16, 28, and 15 cell clusters for low, medium and high confinement. N(MNZ) = 30, 36, 13 cell clusters for low, medium and high confinement. Shaded error bars denote standard error of mean.

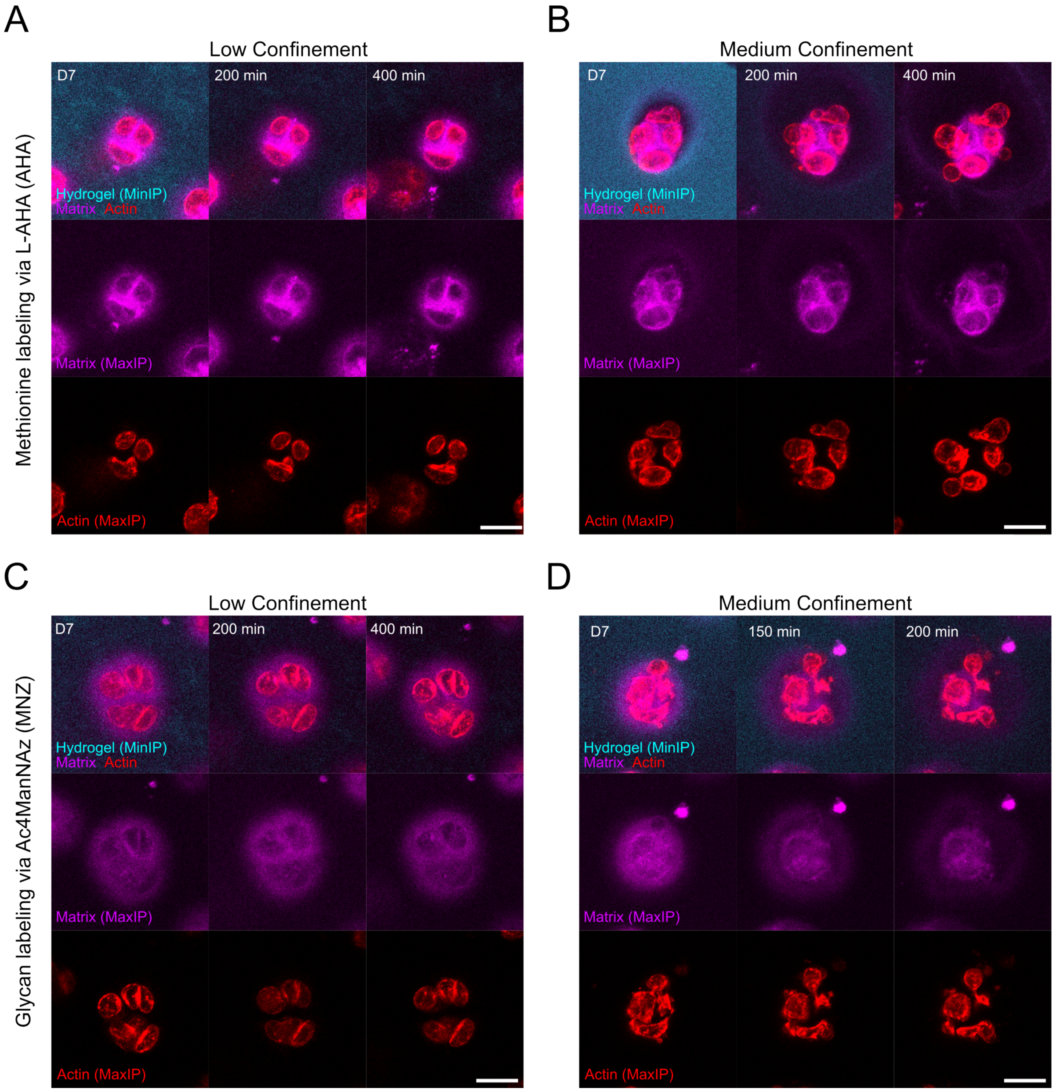

**Figure S7. Additional images of chondrocyte clusters during hydrogel dissolution.** Time-lapse images are obtained on day 7 chondrocyte clusters in low and medium confinement hydrogels, where the hydrogels are dissolved with alginate lyase. A) Low confinement and B) medium confinement hydrogels with fluorescently labelled nascent matrix proteins (labelled with AHA), and C) low confinement and D) medium confinement hydrogels with fluorescently labelled nascent matrix glycans (labelled with MNZ). Note that both AHA and MNZ labelled matrix areas undergo expansion during hydrogel dissolution in medium confinement. Scale bars 25 μm.

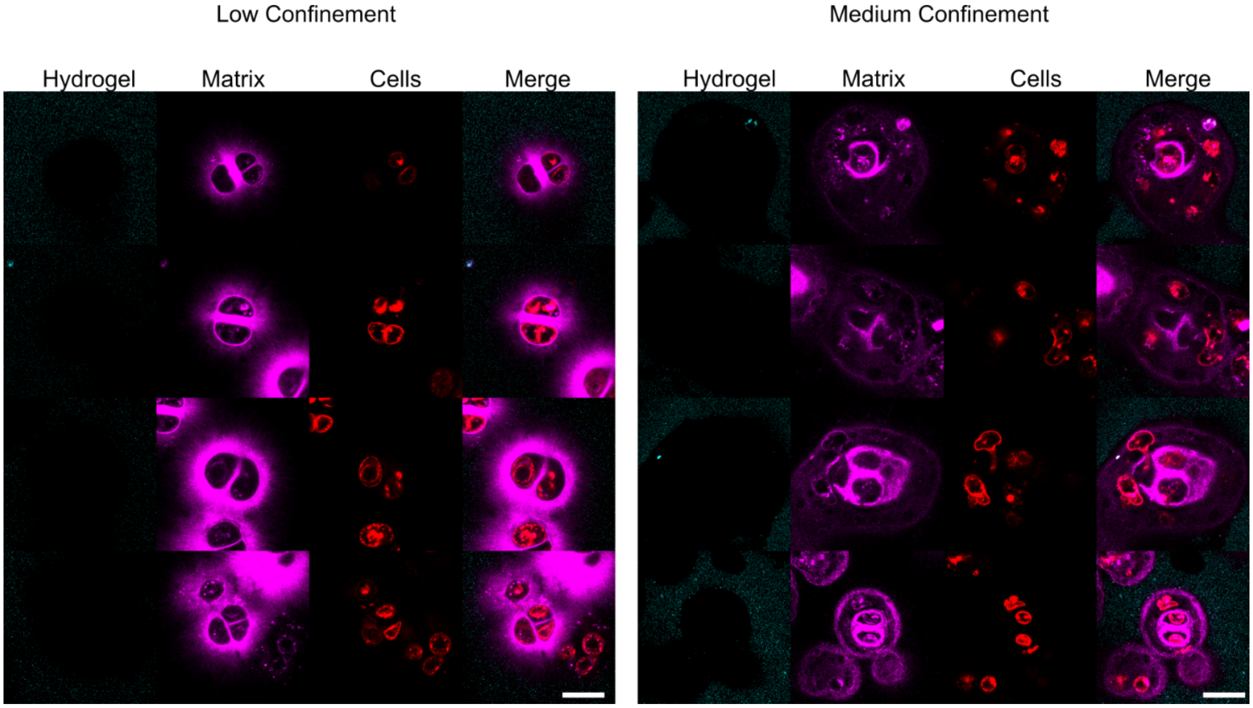

**Figure S8. Post-mortem images of chondrocyte clusters after hydrogel dissolution.** Hydrogels were dissolved using ethylenediaminetetraacetic acid (EDTA) on low confinement and medium confinement hydrogels at day 7, and imaged 24 hours after. Four representative example images are shown per condition. The matrix proteins are labelled with AHA. Scale bars 25 μm.

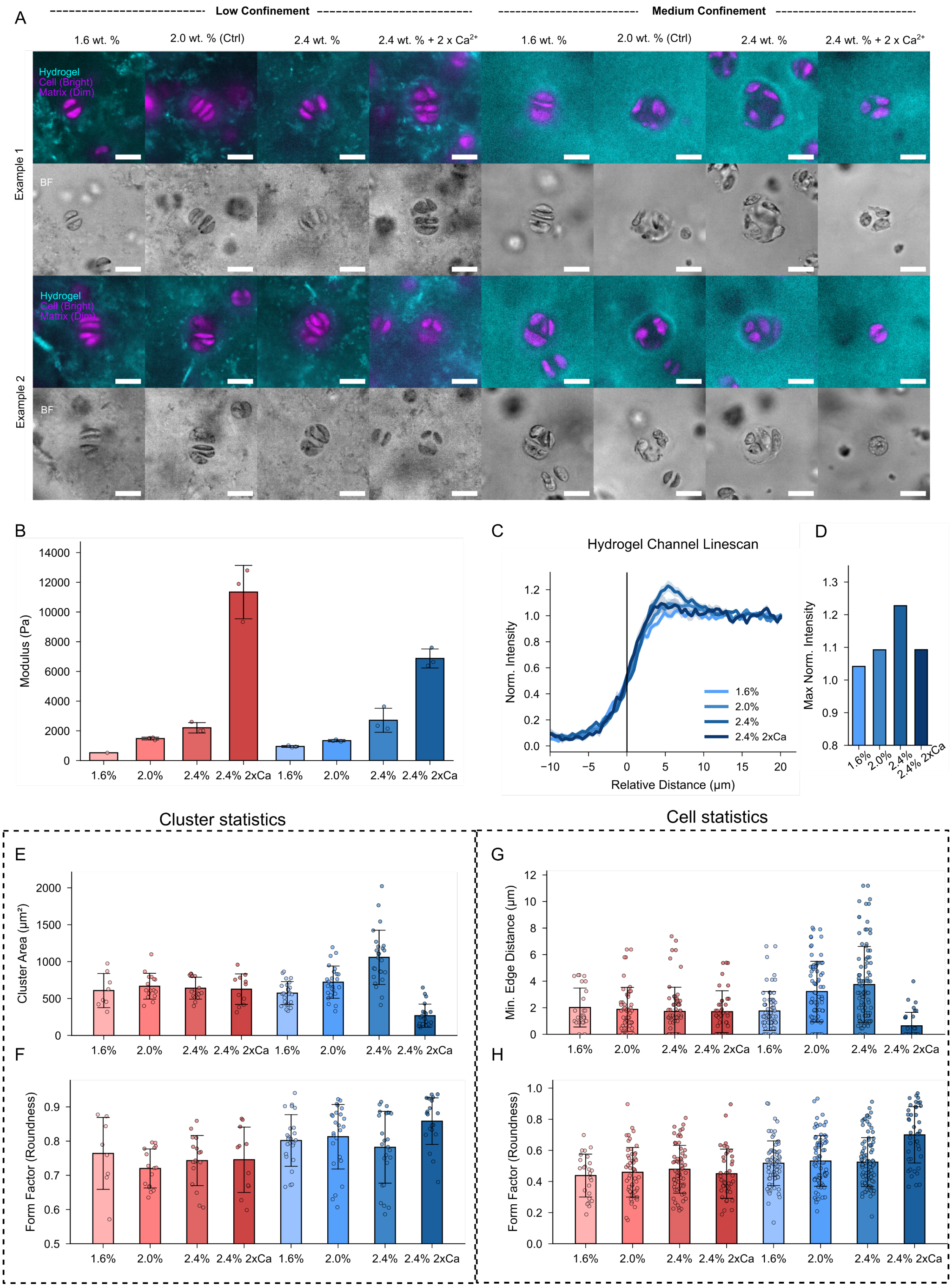
A
Low Confinement
Medium Confinement
1.6 wt. %
2.0 wt. % (Ctrl)
2.4 wt. %
2.4 wt. % + 2 x Ca2+
Hydrogel
Cell (Bright)
Matrix (Dim)
BF
Example 1
Example 2
B
Modulus (Pa)
14000
12000
10000
8000
6000
4000
2000
0
1.6%
2.0%
2.4%
2.4% 2xCa
C
Hydrogel Channel Linescan
Norm. Intensity
Relative Distance (μm)
1.6%
2.0%
2.4%
2.4% 2xCa
D
Max Norm. Intensity
Cluster statistics
Cell statistics
E
Cluster Area (μm²)
F
Form Factor (Roundness)
G
Min. Edge Distance (μm)
H
Form Factor (Roundness)

**Figure S9. Effect of hydrogel elastic moduli on chondrocyte morphology.** Evaluation of the effect of hydrogel elasticity on chondrocyte morphology, where the shear moduli were varied by changing the polymer and calcium cross-linker concentrations (see Supplementary Notes for details). A) Representative day 7 chondrocyte cluster morphology in low and medium confinement hydrogels with varying shear moduli. B) Characterization of the shear moduli. C) Representative intensity linescans of the medium confinement hydrogel channel between the four elasticity conditions, and D) Quantification of the peak of the densified hydrogel channel at the hydrogel wall. E) Cluster area and F) Form factor = $(4\pi\mathrm{Area})/(\mathrm{Perimeter}^2)$ of the cell clusters in low and medium confinement hydrogels of varying shear moduli. G) Minimum cell-cell edge distance and H) form factor of individual cells in the clusters in low and medium confinement hydrogels of varying shear moduli. Note the similarities between the trend of cluster area in medium confinement with different shear moduli, and the cluster areas in low, medium, and high confinement hydrogels characterized in Fig. 3F, as well as the similarities between the trend of minimum cell-cell edge distance in medium confinement with different shear moduli, and minimum cell-cell edge distance in low, medium, and high confinement hydrogels characterized in Fig. 2J. These results highlight how nascent matrix confinement can also be tuned by controlling the modulus of medium confinement hydrogels, allowing the sampling of both low and high hydrogel confinements.

Supplementary Notes: The storage moduli of low and medium confinement hydrogels were adjusted in the following manner. The baseline values for these conditions used throughout the study were 2.0 wt. % polymer and 28 mM Ca (for LC) and 10 mM Ca (for MC). To vary hydrogel elasticity, polymer concentration was rescaled according to the percent values in the figures, with calcium concentration proportionately adjusted as well (e.g. 2.4 % means a 20 % increase in both polymer and calcium concentration). For 2.4 % 2xCa, we instead performed a 100 % increase in calcium concentration, such that we had 56 mM Ca for LC and 20 mM Ca for MC.

For fluorescent imaging of cell clusters in these hydrogels, the hydrogel was first fixed using 4% paraformaldehyde suspended in DMEM for 1 hr at room temperature, after which the paraformaldehyde was rinsed twice in DMEM and finally in TBS with 3 mM $CaCl_2$. DBCO-555 click staining of nascent proteins was performed after this fixation protocol, which led to the staining of intracellular proteins.[4]

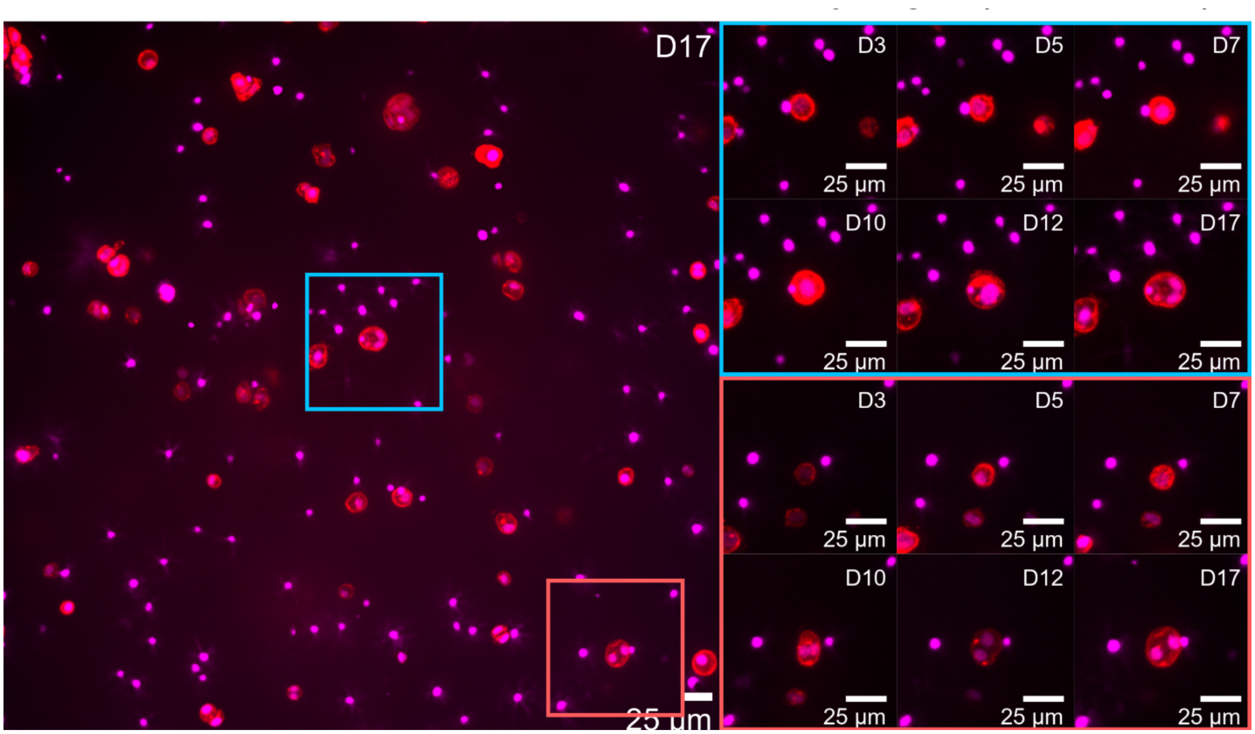

**Figure S10. Additional examples of failed cytokinesis in high confinement hydrogels.** Two examples of failed cytokinesis in high confinement hydrogels occurring in one field-of-view during longitudinal time-lapse imaging. Note the abundance of dead nuclei fragments as well. Red: Cell; Purple: Nucleus. Scale bars 25 µm.

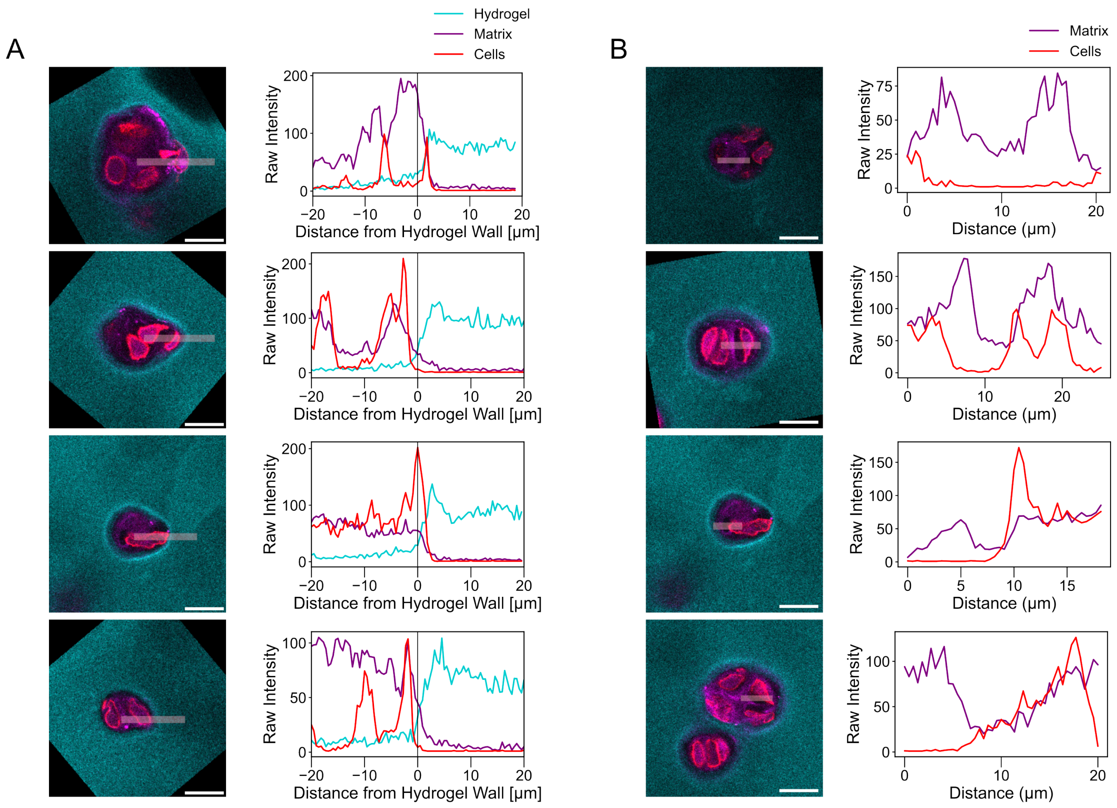

**Figure S11. Chondrocytes leave their nascent matrix cavity to move towards the hydrogel wall.** A) Four representative examples of cells pushing out towards the hydrogel cavity wall, with intensity linescans demonstrating a strong localization of cells near the hydrogel wall. B) Four representative examples of cells leaving holes in the nascent matrix, with intensity linescans demonstrating an absence of cell signals in the nascent matrix cavity. Cyan: Hydrogel, Red: Cell; Purple: Matrix labeled with AHA. Scale bars 25 μm.

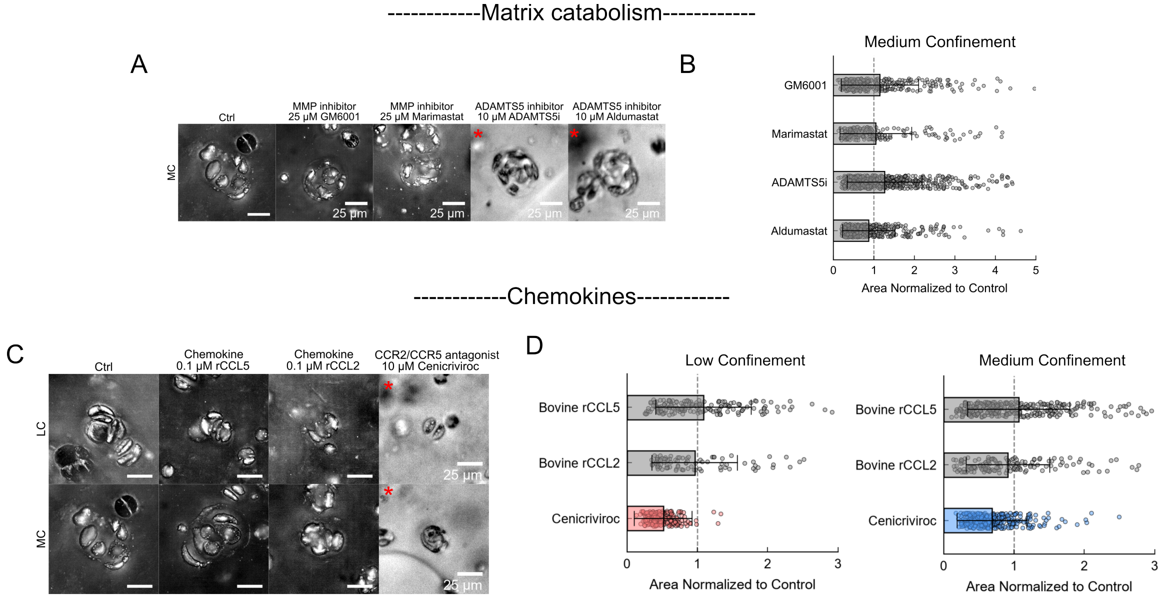

**Figure S12. Perturbation of matrix catabolism and chemokines on chondrocyte cluster morphology.** All pharmacological perturbation studies are done by adding inhibitors to the cell culture media starting from day 1 of culture; control experiments are done by adding a 0.1% DMSO vehicle (except in the case of recombinant CCL2 and CCL5). All imaging experiments are performed on day 7. A) Representative bright-field cluster morphology of chondrocyte clusters in medium confinement hydrogels with broad-spectrum matrix metalloproteinase (MMP) inhibitors GM6001 and Marimastat, and ADAMTS5-specific inhibitors ADAMTS5i (MedChemExpress) and Aldumastat. *denotes images obtained on a 10x objective which gave different phase contrast. Scale bars 25 µm. B) Projected area of clusters in the hydrogels after these perturbation treatments. All areas are characterized by a Cellpose-based automatic segmentation algorithm (see Supplementary Notes). Bars are colored to denote perceptible differences in cluster morphology (grey: visually indistinguishable from control, colored: visually distinguishable from control). C) and D) Same as A and B, but with recombinant CCL5, recombinant CCL2, and CCR2/5 dual antagonist Cenicriviroc in low and medium confinement hydrogels.

Supplementary Notes: Segmentation of cell cluster areas in inhibition studies was done using a custom-trained Cellpose-SAM model fine-tuned on manual annotations of bright-field images of the cell clusters in low and medium confinement hydrogels. Note that, while these trained models were useful for quantifying smaller day 7 clusters here, we found them to be less accurate for larger clusters formed during the 17-day longitudinal study in Fig. 3, for which manual segmentation was preferred.

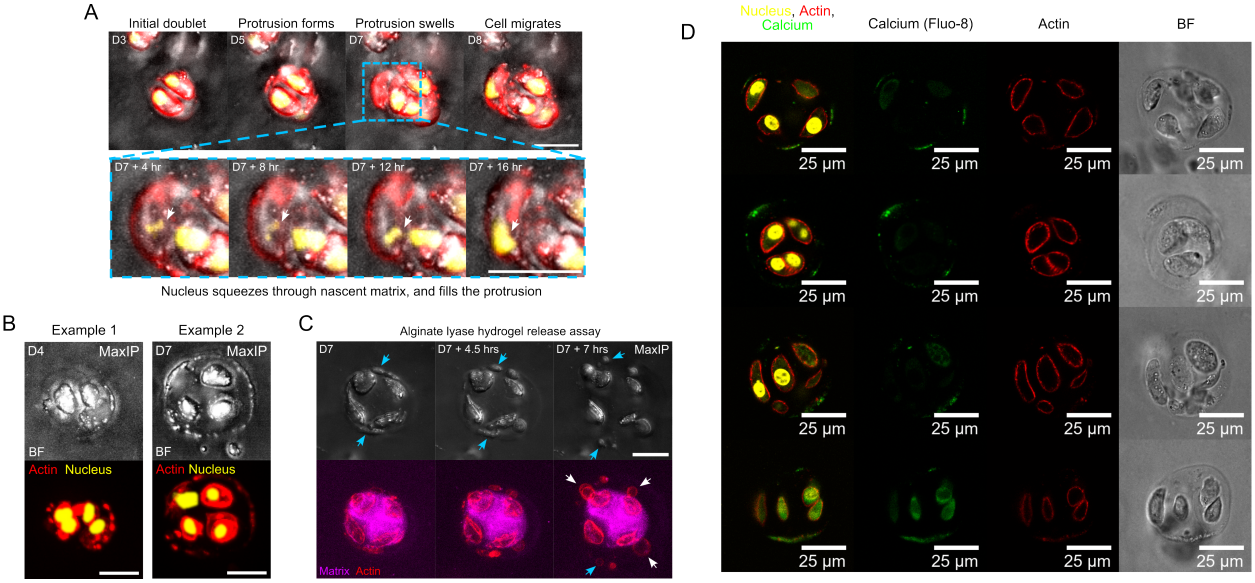

**Figure S13. Nuclear piston and extracellular blebbing during cell migration through the nascent matrix in medium confinement.** A) Supplementary demonstration of the mitosis-driven migration mechanism proposed in Fig. 4H, notably highlighting the squeezing of the nucleus through the narrow pores of the nascent matrix. B) Representative examples of actin-rich strands at the cellular periphery / hydrogel interface. C) Images of cell clusters during hydrogel dissolution via alginate lyase. The actin-rich strands disintegrate into circular blebs (blue arrow). Cells also appear to escape the nascent matrix (white arrow). D) Four example fluorescent images of cell clusters showing enrichment of calcium in the actin-rich strands, co-localized with actin. For this imaging protocol, hydrogels were stained with Hoechst 33342 (1:1000, Invitrogen) and Fluo-8 (5 µM, AAT Bioquest) for 1 hr, and rinsed thrice in DMEM and imaged using a Leica SP8 confocal microscope. Scale bars 25 µm.

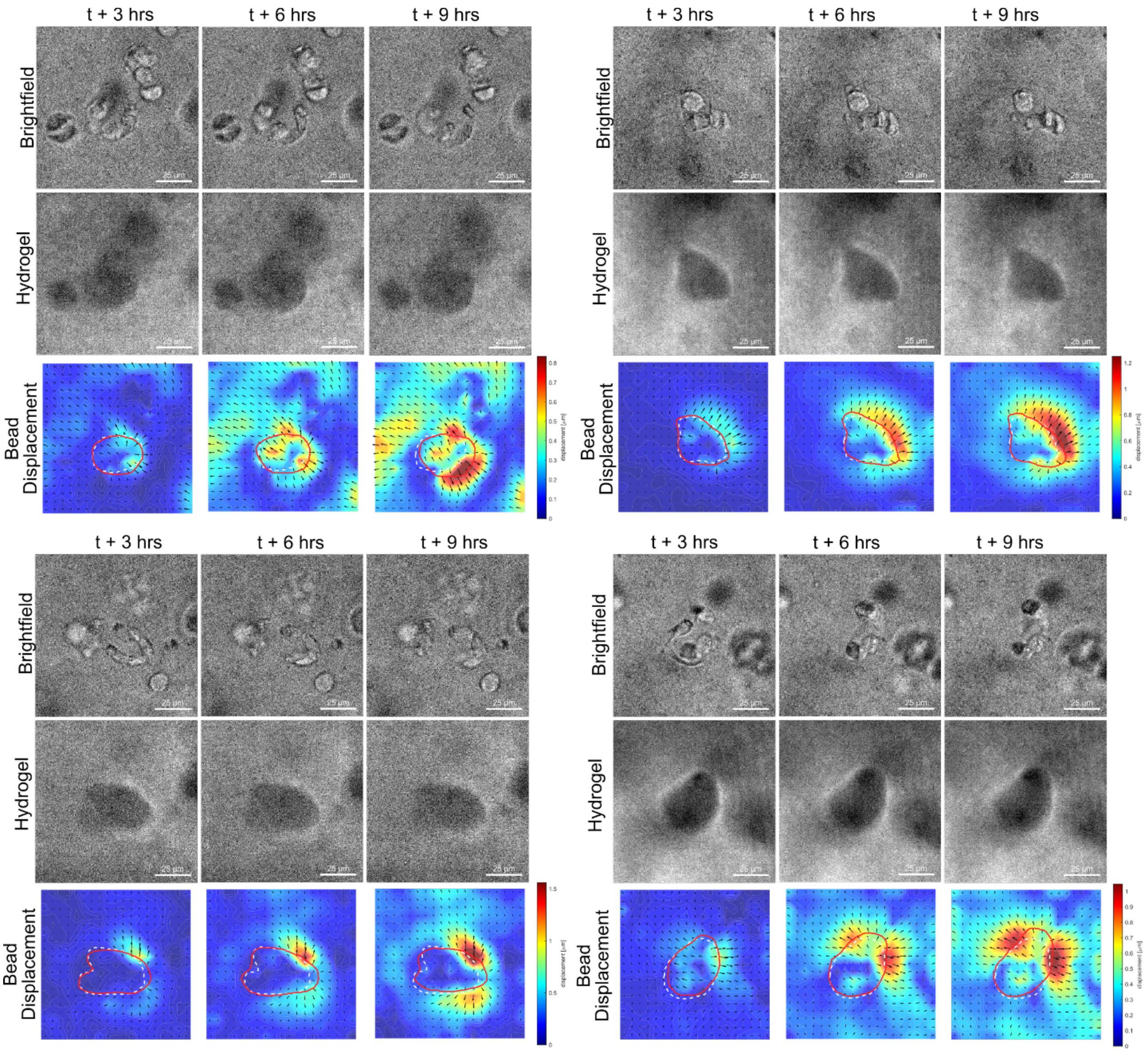

**Figure S14. Bead displacement characterization of medium confinement hydrogels during cell migration.** Dashed outline on the PIV panel indicates initial cell cluster region, and solid outline indicates final cell cluster region. Four representative examples at day 7 are shown. Scale bars 25 µm.

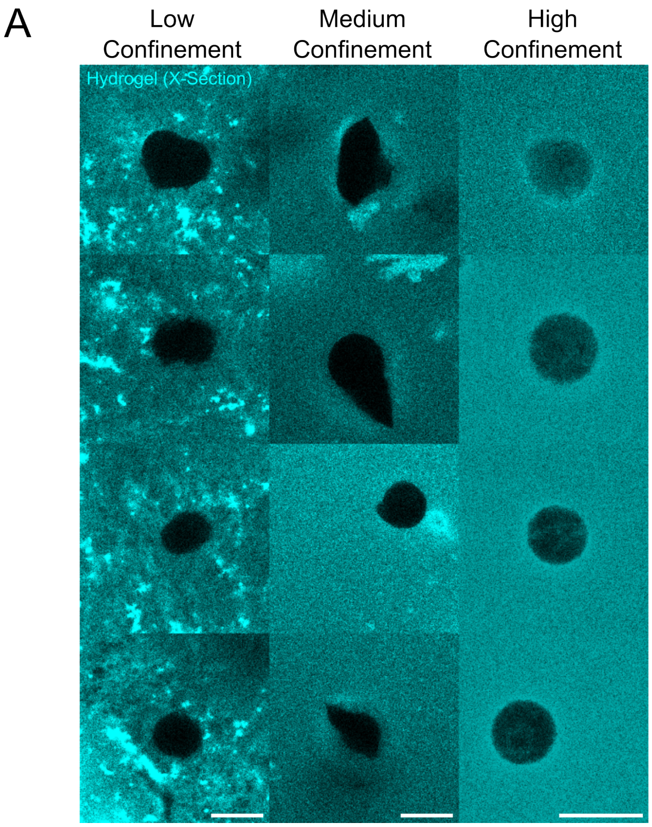

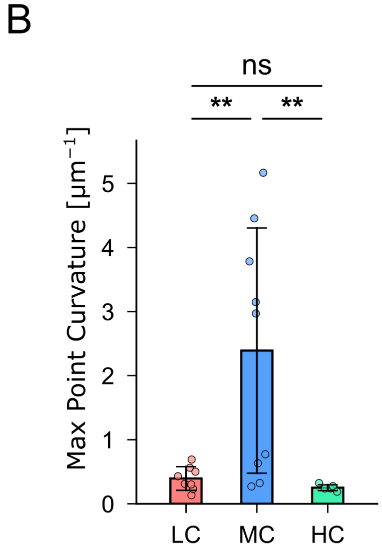

**Figure S15. Chondrocyte migration leads to cavity fracture in medium confinement.** A) Images of the hydrogel cavities after nascent matrix dissolution. Collagenase was used to dissolve the nascent matrix at day 7, and imaging was done after 24 hours. Four representative examples are shown. Scale bars 25 μm. B) Quantification of the maximum point curvature of the cavities, done via the Kappa plugin in ImageJ.

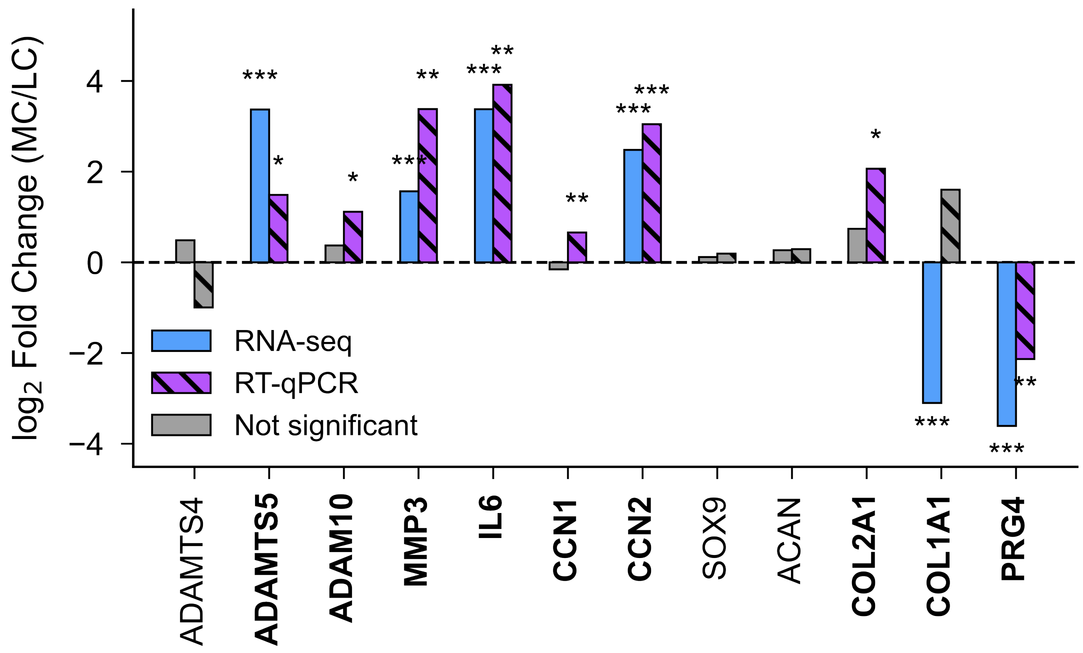

**Figure S16. Cross-validation between RNA-seq and RT-qPCR.** Comparison of relative log2 fold change between MC and LC of key chondrocyte genes analyzed by RNA-seq and RT-qPCR at day 7. Bolded genes denote genes where at least one measurement technique revealed a statistically significant difference. Asterisks denote statistical significance within each measurement.
