## Supplementary Tables for "Hydrogel confinement of the secreted nascent matrix imposes solid stress and mediates pathology of chondrocytes"

**Table S1. Reagents**

| Reagent | Source | Category |
|---|---|---|
| Dulbecco's Phosphate-Buffered Saline (DPBS) | Gibco | General / Cell culture |
| Low glucose Dulbecco's Modified Eagles Medium (DMEM) | Gibco | |
| Dimethyl Sulfoxide (DMSO) | Medchemexpress | |
| 1M HEPES buffer | Gibco | |
| 100X Minimum Essential Medium Non-Essential Amino Acids (MEM-NEAA) | Gibco | |
| 100X Antibiotic-Antimycotic (Anti-Anti) | Gibco | |
| L-Ascorbic Acid-2-Phosphate Sesquimagnesium Salt Hydrate | Sigma | |
| Standard Fetal Bovine Serum (FBS) | Cytiva | |
| Calcium sulfate | Sigma | |
| Calcium chloride dihydrate | Sigma | |
| Alginate lyase | Sigma | |
| Ethylene diamine tetraacetic acid (EDTA) | Sigma | |
| Sigmacote | Sigma | |
| Low gelling temperature agarose | Thermo Scientific | |
| Bovine serum albumen (BSA) | Sigma | |
| Tris-buffered saline (TBS) | Thermo Scientific | |
| Mineral oil BioReagent | Sigma | |
| Trypsin-EDTA (0.05%) | Gibco | |
| 500X Gentamicin-Amphotericin | Gibco | |
| Collagenase 2 | Worthington | |
| Collagenase 4 | Worthington | |
| Pronova UP VLVG alginate (Low molecular weight alginate, less than 75 kDa) | Novamatrix | Polymer reagents |
| Pronatal LF 20/40 alginate (High molecular weight alginate, greater than 200 kDa) | FMC Biopolymer | |
| 8-arm PEG Norbornene 20 kDa | Jenkem | |
| 2-arm PEG Thiol 5 kDa | Jenkem | |
| FITC-PEG Thiol 1 kDa | Creative PEGWorks | |
| Lithium phenyl-2,4,6-trimethylbenzoylphosphinate (LAP) | Sigma | Alginate synthesis |
| N-(3-Dimethylaminopropyl)-N'-ethylcarbodiimide hydrochloride | Sigma | |
| Sulfo-N-hydroxysulfosuccinimide sodium salt | Bioworld | |
| 6-aminofluorescein | Sigma | |
| Hydroxylamine hydrochloride | Sigma | |
| 4-Morpholineethanesulfonic acid (MES) buffer | Sigma | |
| Dialysis tubing | Spectrum Laboratories | |
| Activated charcoal | Sigma | |

| | | |
|---|---|---|
| High glucose DMEM, no glutamine, methionine, or cystine | Gibco | Metabolic labeling |
| 200 mM L-Glutamine | Gibco | |
| 100 mM sodium pyruvate | Gibco | |
| L-Methionine | Sigma | |
| L-Cystine | Thermo Scientific | |
| L-Azidohomoalanine | Vector Labs | |
| N-azidoacetylmannosamine-tetraacetylated | Vector Labs | |
| N-azidoacetylgalactosamine-tetraacetylated | Vector Labs | |
| AZDye 488 / 555 DBCO | Vector Labs | Fluorescent imaging |
| SPY555-DNA | Spirochrome | |
| SiR-Actin | Spirochrome | |
| Calcein-AM | Invitrogen | |
| Ethidium homodimer-1 | Invitrogen | |
| Fluosphere carboxylated microspheres (0.2 um, 540/560 nm) (F8809) | Invitrogen | |
| Sodium acetate trihydrate | Sigma | Hydroxyproline assay |
| Sodium citrate dihydrate | Sigma | |
| Citric acid monohydrate | Sigma | |
| Chloramine T trihydrate | Sigma | |
| Perchloric acid | Sigma | |
| p-dimethylaminobenzyldehyde (DMAB) | Sigma | |
| Papain from papaya latex | Sigma | |
| L-cysteine | Sigma | |
| GM6001 | Medchemexpress | Inhibition study |
| Marimastat | Medchemexpress | |
| ADAMTS5 inhibitor (ADAMTS5i) | Medchemexpress | |
| Aldumastat | Medchemexpress | |
| Bovine rCCL2 | Kingfisher biotech | |
| Bovine rCCL5 | Kingfisher biotech | |
| Cenicriviroc | Medchemexpress | |
| Genomic lysis buffer | Zymo Research | DNA/RNA assay |
| RNA lysis buffer | Zymo Research | |
| DNA/RNA shield | Zymo Research | |
| Quick-RNA microprep kit | Zymo Research | |
| Primescript RT reagent kit | Takara Bio | |
| Power SYBR-Green master mix | Applied Bio | |
| Qubit 1x dsDNA br assay kit | Invitrogen | |

### Table S2. Primers

| Gene | Forward primer (5′→3′) | Reverse primer (5′→3′) | Ref. |
|---|---|---|---|
| CCN1 | TTACGCTGGATGTTCGAGTG | AACCGCATCTTCACAGTCCT | 1 |
| CCN2 | CAAGCAGCTGAGCGAGTTGT | TGGTACACAGTTCCTCCGAAA | 2 |
| ADAMTS5 | AGCATTGACGCATCCAAACCCT | TTCGTGGTAGGTCCAGCAAACA | 3 |
| P16 | CCCTCGTGCTGATGGCTAGT | CCCATCATCATCACCTGGTCTA | 4 |
| P21 | CGCCAGCTGAGGTGTGAG | ATGGCACCTGTGGCTCTTCT | 4 |
| P53 | ACTCTTCAGATCCGTGGGTTTA | CCATCCAGAGCATCCTTCAG | 4 |
| IL6 | TGAGTGTGAAAGCAGCAAGG | AGCAAATCGCCTGATTGAAC | 5 |
| PRG4 | AGAAAACCCGATGGCTATGA | TCGCCCATCAGTCTAAGGAC | 6 |
| MMP13 | TGGTCCAGGAGATGAAGACC | TGGCATCAAGGGATAAGGAA | 7 |
| ADAMTS4 | AACTCGAAGCAATGCACTGGT | TGCCCGAAGCCATTGTCTA | 7 |
| IL1B | CCACCCAGGGATCCTATTCT | TCATTTCGTTGATGGGTTCA | 7 |
| SOX9 | ACGCGGCCCCAGGAGAAC | CGGATGCACACGGGGAACTT | 8 |
| COL2A1 | GTGGGGCAAGACTATGATCG | TGCAATGGATTGTGTTGGTT | 8 |
| COL1A1 | ACATTGGCCCAGTCTGTTTC | GGGAGGGGGAGTGAATTAAA | 8 |
| ACAN | CACCACAGCAGGTGAACTAGA | GCTTGCTCCTCCACTAATGTC | 8 |
| GAPDH | CCTTCATTGACCTTCACTACATGGTCTA | TGGAAGATGGTGATGGCCTTTCCATTG | 9 |
| MMP3 | TGGACCTGGAAAAGTTTTGG | AGCTTCACGTTCGGTTGAGT | 10 |
| ADAM10 | CATCTGGGGACAAACTTAACAACA | CCCATTTCCACAAATAGGTTGGC | 11 |
